# ADBP-1 regulates ADR-2 nuclear localization to control editing substrate selection

**DOI:** 10.1101/2023.05.14.540679

**Authors:** Berta Eliad, Noa Schneider, Orna Ben-Naim Zgayer, Yarden Amichan, Fabian Glaser, Emily A. Erdmann, Suba Rajendren, Heather A. Hundley, Ayelet T. Lamm

**Author notes:** Corresponding author: Ayelet T. Lamm. Faculty of Biology, Technion - Israel Institute of Technology, Technion City, Haifa 3200003, Israel. These authors contributed equally to this work (Berta Eliad, Noa Schneider).

## Abstract

Adenosine-to-inosine (A-to-I) RNA editing, catalyzed by ADAR enzymes, is a prevalent and conserved RNA modification. While A-to-I RNA editing is essential in mammals, in *Caenorhabditis elegans*, it is not, making them invaluable for RNA editing research. In *C. elegans*, ADR-2 is the sole catalytic A-to-I editing enzyme, and ADR-1 is an RNA editing regulator. ADAR localization is well-studied in humans but not well-established in *C. elegans*. In this study, we examine the cellular and tissue-specific localization of ADR-2. We show that while ADR-2 is present in most cells in the embryo, at later developmental stages, its expression is both tissue- and cell-type-specific. Additionally, both ADARs are mainly in the nucleus. ADR-2 is adjacent to the chromosomes during the cell cycle. We show that the nuclear localization of endogenous ADR-2 depends on ADBP-1, not ADR-1. In *adbp-1* mutant worms, ADR-2 is mislocalized, while ADR-1 is not, leading to decreased editing levels and *de-novo* editing, mostly in exons, suggesting that ADR-2 is also functional in the cytoplasm. Besides, mutated ADBP-1 affects gene expression. Furthermore, we show that ADR-2 targets adenosines with different surrounding nucleotides in exons and introns. Our findings indicate that ADR-2 cellular localization is highly regulated and affects its function.

## INTRODUCTION

RNA editing is a common post-transcriptional process essential for RNA function (1). The most prevalent type of RNA editing is A-to-I RNA editing (2). In this process, adenosine (A) within double-stranded RNA is deaminated into inosine (I), which is recognized by the splicing and translational machinery as guanosine (G) (3, 4). The enzymes that catalyze this conversion are called Adenosine Deaminases Acting on RNA (ADAR) (5). This seemingly minor modification has an extensive effect; inosine, a non-canonical nucleotide, signals the cell that the modified RNA is not foreign, preventing undesirable activation of the immune system (6). RNA editing can also alter proteins’ amino acid sequence and generate protein isoforms (7). Moreover, alteration of editing occurs in many neurological disorders and different cancers (8–13).

In humans, two catalytically active ADARs are known: ADAR1 and ADAR2; both are essential, expressed in most tissues, and target double-stranded RNA and DNA/RNA hybrids (4, 14–17). ADAR1 has two isoforms capable of shuttling between the nucleus and cytoplasm (18). ADAR1 p110 is constitutively expressed and mostly nuclear, while ADAR1 p150 is interferon-induced and thus could be expressed, for example, during viral infection (18). p150 tends to accumulate in the cytoplasm owing to a nuclear export signal (NES) found in its unique N-terminal region. A non-classical nuclear localization signal (NLS) overlaps the third double-stranded RNA binding domain (dsRBD) in both isoforms (19–21). The localization of the different isoforms is thought to be one of the regulatory mechanisms affecting ADAR1 RNA editing activity (19, 22, 23). Although ADAR2 is nuclear, it can be sequestered into the nucleolus via an NLS found in its dsRBD (also in the N-terminus region of the enzyme) (24). ADARs have been shown to be essential in mammals; for example, mice containing a homozygous deletion for ADAR2 die shortly after birth (25).

While ADAR1 edits mainly non-coding regions, ADAR2 edits non-coding and coding regions that can lead to protein recoding (18, 25). Previous works tried to find if ADAR enzymes have preferable nucleotides surrounding their target site (27–33). Although each work done on human ADARs showed slightly different motifs (27–30), they found in common that human ADAR1 and ADAR2 prefer uridine at the 5’ of the edited sites and guanosine at their 3’.

In contrast to humans, in *Caenorhabditis elegans,* deletion of either or both of its ADAR genes is not lethal (34), making this well-characterized model organism ideal for researching the function of RNA editing (35). *C. elegans* has two ADAR genes: ADR-1 and ADR-2 (35). The function of ADR-1 is mainly regulatory (36–39), while ADR-2 is catalytically active and edits mainly non-coding sequences (34, 36, 40). Previous work showed that ADR-1 protein was primarily located in the nuclei and a smaller fraction in the cytoplasm (41). In contrast to ADR-1, previous studies have been unsuccessful at determining the expression pattern of the ADR-2 protein by using translational reporters, presumably because *adr-2* is situated in a six-gene operon and there are undefined control areas, or it is possible that overexpression of *adr-2* is lethal (34).

The first indication of regulation of ADR-2 localization came from the finding that ADBP-1 (ADR-2 Binding Protein 1) was shown to affect the subcellular localization of a heterologous expressed ADR-2 transgene (42), causing it to be localized also in the cytoplasm instead of only in the nucleus. ADBP-1 was shown to interact with ADR-2, and in the absence of functional ADBP-1, four known targets of ADR-2 did not undergo RNA editing (42). However, ADBP-1’s effect on the endogenous ADR-2 was not determined.

To gain more insight into the localization of ADR-2 and, thus, the mechanisms of regulation of its RNA editing activity, we performed immunofluorescence studies and RNA-seq experiments. We found that ADR-2 is ubiquitously expressed in wild-type embryos and adjacent to the chromosomes throughout the cell cycle. In contrast, except in the gonads, ADR-2 is not ubiquitously expressed in the worm’s tissues at later developmental stages or in the sperm. We show that both ADR-1 and ADR-2 are localized in the nucleus. Without a functional ADBP-1 protein, ADR-2 appears cytoplasmic, while ADR-1 remains mainly in the nucleus. Mislocalized ADR-2 can still edit mRNA; however, this editing happens at lower levels than when ADR-2 is localized primarily in the nucleus. Although editing levels decrease, ADR-2 mislocalization causes *de-novo* editing, which appears to be sporadic. Additionally, we found that the nucleotide signature surrounding adenosines targeted by ADR-2 differs between untranslated regions, coding exons, and introns but is strain-independent. Our results suggest that the localization of ADR-2 is highly regulated and likely affects the editing levels and expression of cellular transcripts.

## MATERIALS AND METHODS

### Maintenance and handling of *C. elegans* strains

Experiments were performed with the wild-type Bristol strain N2(43), BB21 (*adr-1(tm668)* I; *adr-2(ok735)* III (34)), RB886 (*adr-2(ok735)* III (41)), QD1 (*adbp-1(qj1)* II(42)), HAH36 (V5:adr-1;FLAG:adr-2), ALM132 (V5:*adr-1;adbp-1(qj1) II*), and ALM517 *(adr-1(uu49*). ALM517 was obtained by outcrossing 3 times and separating *adr-1(uu49)* from strain BB239 (*adr-1(uu49)* I; *adr-2(uu28)* III) (44). HAH36 (V5:adr-1;FLAG:adr-2) (45) was crossed first with N2 to obtain homozygous worms carrying V5:adr-1 only and then with QD1 (*adbp-1(qj1)* II) (42) strain to create strain ALM132 (V5:*adr-1;adbp-1(qj1) II*). All *C. elegans* strains were grown at 15°C on NGM agar 5-cm plates and seeded with E. coli OP50 bacteria. For mRNA-seq libraries preparation, strains were grown at 20°C.

### Rescue strain preparation

The *adbp-1*:g*fp* “L” plasmid (a kind gift from Prof. Manabi Fujiwara (42)) was co-injected with the *rol-6* marker. An injection mix contained 70 ng/µl *rol-6* co-injection marker, 20ng/µl of 1Kb DNA ladder (Thermo Scientific), and 10ng/µl of the *adbp-1*:*gfp* “L” plasmid for a final concentration of 100ng/µl. Worm DNA was checked by PCR for the presence of the *adbp-1*:*gfp* “L” plasmid using AAAAGCTGAAGAAACAGGAC and TTAACATCACCATCTAATTCAAC primers to *adbp-1*.

### DNA and RNA Sanger sequencing

DNA extraction was performed according to a protocol in which worm lysis buffer is applied to a sample of ∼5 worms, followed by cycles of cooling and heating temperatures (−80°C for 15 minutes, 60°C for 60 minutes, 95°C for 15 minutes, finally cooling down to 4°C) (46).

To obtain cDNA, extracted RNA was treated with turbo DNase (Ambion). Then, a reverse transcriptase reaction (Quanta - qScript Flex cDNA Kit) was done using oligo-dT. The amplification products were sequenced by Sanger sequencing.

### mRNA-seq libraries preparation

Worms were washed with M9 and treated with sodium hypochlorite. The embryos were either resuspended in EN buffer and frozen in liquid nitrogen or resuspended in M9 buffer and left overnight in a rotator at 20°C. The hatched and synchronized L1 larvae were placed on NGM media with OP50 until they reached the L4 stage. The L4 larvae were washed with EN buffer and frozen with liquid nitrogen. Frozen pellets were ground to powder with liquid nitrogen chilled mortar and pestle. RNA was extracted by using Direct-zol RNA MiniPrep Plus (ZYMO). The RNA was treated with TURBO DNAse (Ambion). mRNA sequencing libraries were prepared using a TruSeq RNA kit from Illumina and sequenced by Illumina HiSeq 2500.

### Detection of A-to-I RNA editing in RNA-seq

The current A-to-I RNA editing detection is based on at least three different biological RNA-seq replicas of the following strains: N2 (wild-type), ADAR mutant worms (BB21), and *adbp-1* mutant worms (QD1), at the embryo and L4 stages (GSE83133, GSE230883).

The quality of RNA-seq reads was estimated by FastQC (47). Afterward, reads with poor base quality at their edges were trimmed by an in-house script to improve alignment and editing detection. Identical reads were collapsed in all the examined samples by an in-house script. Our in-house scripts can be found at: https://github.com/Lammlab/ADR-2_localization. The reads were aligned to *C. elegans* WS220 genome using Bowtie (48) by the following command: bowtie -p 23 -n 3 -e 120 -a --strata –best --sam -m 2 –un. SAM format files, the alignment output files, were processed to BAM files using SAMtools (49). BAM files of samples representing the same strain and development stage were merged and processed to a pileup format using SAMtools (49). To detect editing of known editing sites (36, 40) in wild-type and *adbp-1* mutant worms, we removed sites that appeared to undergo editing above 3% in the ADARs mutant (BB21 strain) from the pileup files to avoid artifacts that might be derived from non-A-to-I-RNA-editing events. Next, we filtered the base calls of reads aligned to a reference sequence in pileup files so only sites previously identified as being edited remain. In determining the editing levels on those sites, we considered only nucleotide changes with Phred quality ≥ 25 to increase editing detection precision. A site was considered edited if A-to-G or T-to-C (the revered strand) changes appeared to be ≥ 1%, and no more than 1% of other nucleotides were changed along the nucleotides covering the site. The known A-to-I editing sites analysis included a comparison of editing both on a single site level and at the whole gene editing level. To calculate editing at the gene level, we merged bases from aligned reads that belong to the same gene. The final lists of known editing sites that appeared in the used samples are presented in the supplemental tables. The results were plotted using the ggplot2 R package.

The *de-novo* A-to-I RNA editing sites search is based on the pipeline published by Goldstein et al., 2017 (40). However, our study considers the Phred quality of the reads. This analysis included a different pileup file for each sample, and pileup files represented merged data by strain and developmental stage. To identify nucleotide changes in the RNA that are not a result of RNA editing events, we eliminated nucleotide changes derived from the DNA sequence from the pileup files using DNA-seq (40) and single nucleotide polymorphisms dataset (50). Next, we eliminated each nucleotide change in the ADAR mutant from the pileup files. Next, we defined a site as edited if it meets the following criteria: Nucleotide change was considered if its Phred quality ≥ 25. The most abundant nucleotide change among the site’s reads appears in at least two expressed reads. The percentage of the most abundant nucleotide change ≥ 5% of the total expressed reads covering the site. The not abundant nucleotide changes appear to be ≤ 1% of the total expressed reads covering the site. The edited site appeared in at least two biological replicas.

We defined an A-to-I editing site in the *adbp-1* mutant as a *de-novo* if it does not appear to be edited in wild-type samples in the known editing site analysis and the *de-novo* A-to-I RNA editing sites search. To annotate each detected editing site, we used the WormBase ParaSite ws220 annotations database (51, 52).

### Minimum free energy calculation of RNA secondary structures

To calculate the free energy of secondary structures surrounding the detected editing sites or random adenosines, we extracted the 50 nucleotides surrounding the editing sites or the random adenosine from both sides to obtain a sequence of 101 nucleotides. The same was performed for T-to-C nucleotide changes, using their reversed and complement sequences. We used edited and random adenosines located in exons, so sequences originating from unspliced and spliced forms were obtained. Genome reference ws220 and transcriptome reference ws220 were used to extract the unspliced and spliced sequences, respectively. To predict the minimum free energy of the secondary structures, we used the seqfold python package (https://pypi.org/project/seqfold/). Outliers with minimum free energy above 100 were removed from the analysis. We used the Welch two-sample T-test to test if a difference in mean minimum free energy between groups is statistically significant.

### Nearest neighbors surrounding editing sites

We used Logomaker, a Python package (53), to create the logos of the nucleotides surrounding editing sites and random adenosines. The logo represents a probability matrix, which means the probability of observing each possible nucleotide at each possible position within a particular sequence type. The following equation calculated the probability: 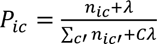, where *P*_*ic*_ represents matrix elements. *C* is the number of possible characters, and λ is a user-defined pseudo count. A probability logo has heights given by these *P*_*ic*_ values.

### Gene expression analysis

The RNA-seq data for A-to-I RNA editing site detection were analyzed for gene expression. The reads were aligned to *C. elegans* WS220 transcriptome using Bowtie (48) using a command that allows multiple alignments for isoform-aware alignments: bowtie -p 3 -n 3 -e 120 -a --strata --best --sam -m 10 –un. We pre-filtered the count genes, so only genes with at least ten reads with a coefficient of variation < 1 were analyzed. To test the significance of lncRNA and 3’UTR edited genes to all the expressed genes in all samples, we applied the Welch two-sample t-test between the chosen two groups (two-sided). We used DESeq2 (54), an R package, to analyze gene counts and identify differentially expressed transcripts. We considered genes as differentially expressed (DE) between the wild-type and mutated strains that adhered to the following criteria: |log2FoldChange| > 1 and padj < 0.05 at the embryo stage and |log2FoldChange| > 2 and padj < 0.05 at the L4 stage. The results are exhibited by volcano plots created by the ‘EnhancedVolcano’ package in R (55). We applied enrichment analysis of the DE genes by avoiding the mutated genes in WormBase Enrichment Suite (56, 57). P-values in Venn diagrams were determined by hypergeometric distribution using the P_hyper_ function in R.

### Immunostaining

In order to visualize *C. elegans* embryos, adult worms containing embryos were fixed and prepared for immunostaining according to a previously described fixation protocol with methanol-acetone (58). In order to visualize larvae and adult worms, a mix of worms present at different stages was fixated with 1% formaldehyde and permeabilized as described previously (59). Primary rabbit anti-ADR-2 (IU529) (38) was used at a 1:50 dilution. Donkey anti-rabbit Secondary Antibody, Alexa Fluor 568 (Life Technologies, #A10042), was used at a 1:200 dilution. Primary mouse anti-MH27 (# MH-27-s) from DSHB (Developmental Studies Hybridoma Bank, The University of Iowa), was used at a 1:300 dilution (gift from Benjamin Podbilewicz’s lab) with secondary antibody Alexa Fluor 488 donkey anti-mouse (Life technologies, #A21202) at a 1:500 dilution. Primary mouse anti-V5 (Invitrogen) or rabbit anti-V5 (Signaling Technology) were used at a 1:500 dilution with secondary antibody Alexa Fluor 647 goat anti-mouse (Jackson immunoresearch) or Alexa Fluor 488 Goat anti-rabbit (Invitrogen) diluted to 1:100, respectively. Primary rabbit anti-Lem-2 (Novus) was used at a 1:1000 dilution with secondary antibody Alexa Fluor 488 Goat anti-rabbit (Alexa Mol Probes) at a 1:500 dilution. Primary mouse anti-Flag (Sigma) was used at a 1:750 dilution with secondary antibody Alexa Fluor 647 goat anti-mouse (Jackson immunoresearch) diluted to 1:100. DAPI (4′,6-diamidino-2-phenylindole, Sigma) was used at a 1:1000-1:2000 dilution for DNA staining.

Microscope images of embryos and most of the adult worms were obtained with a Spinning Disk Confocal microscope from Nikon with CSU-W1 Confocal Scanner Unit with dual camera from Yokogawa. The objective used for all images was x100 oil (NA = 1.45) CFI PLAN APOCHROMAT. For the embryo images exposure was set at 200 milliseconds for both channels (405nm for DAPI and 561nm for ADR-2 antibody). For the adult and larva images exposure was set at 400 milliseconds for 405nm and 100 milliseconds for both 561nm and 488nm (for MH27 antibody). Laser intensities were set at 35% for 405nm, 22.1% for 568nm and 15% for 488nm. Z-stacks for all images were obtained in a range of 7 μm in the z-axis. The acquisition was performed using the NIS-Elements AR software. Maximal intensity z-projection of the stacks was created using Fiji (imageJ, NIH) for images of the hermaphrodite body and tail belonging to wild-type and *adbp-1^−/−^* strains. The rest of the images are individual slices taken from their corresponding z-stacks (hermaphrodite head for both strains, embryo and cell cycle images).

Images of larvae and adults were also obtained using an inverted microscope (Nikon Ti ECLISPE) with a Confocal Spinning Disk (Yokagawa CSU-X), with an Andor iXon3 camera (DU-897-CSO-#BV). The objectives used were a 60x oil Plan Apochromat (NA = 1.4) lens or a 40x oil Plan Fluor (NA= 1.3) lens. Some images were enhanced with an additional x1.5 amplification. The laser intensity was set at 7%, exposure at 100 milliseconds and the gain at 300 for both channels (405nm for DAPI and 561nm for ADR-2 antibody). Z-stacks were obtained in a range of 5 μm in the z-axis. The software used was the Molecular Devices Metamorph.

All microscope images were corrected for brightness and contrast together with the controls, merged, and stacked to RGB using Fiji (imageJ, NIH).

Quantifying signal intensity was done using the ImageJ measurement tool (60). The area was selected for each cell measured in the nucleus, cytoplasm, and background outside the worm. The position of the nucleus was determined by using DAPI staining. Nucleus/cytoplasm expression fold change was determined after subtracting the background mean grey value from both nucleus and cytoplasm values. We used the Mann-Whitney U-Test to calculate the statistical significance of the nucleus/cytoplasm fold change between different strains.

### Immunoprecipitation

Mixed-stage worms were washed with IP buffer (50 mM HEPES [pH 7.4], 70 mM K-Acetate, 5 mM Mg-Acetate, 0.05% NP-40 and 10% glycerol) and frozen at −80°C. Frozen worm pellets were ground with a cold mortar and a pestle on dry ice. The cell lysate was centrifuged at maximum speed to remove cellular debris. Protein concentration was measured with Bradford reagent (Sigma-Aldrich), and 5 mg of the worm lysate was added to anti-Rabbit IgG magnetic Dynabeads (Fisher) coated with the same custom ADR-2 antibody described above. After incubation for 1 hour on the cold room rotator, protein-bound beads were washed with a wash buffer (0.5 M NaCl, 160 mM Tris-HCl [pH 7.5]). A portion of the IP (1/10) was stored in an SDS loading buffer and used for immunoblotting (Supplemental Figure 14). The remaining beads were stored with 100 ml of 1X TBS (0.11 M NaCl, 16 mM Tris-HCl [pH 7.5]) and flash frozen at −80°C.

### Mass spectrometry and data analysis

The beads containing immunoprecipitated ADR-2 and other interacting proteins were subjected to proteolytic digestion and LC-MS/MS at the Indiana University School of Medicine Proteomics core. The mass spectrometry results were analyzed using Scaffold4 (Version 4.8.9), and the statistical significance between the samples was calculated using Fisher’s t-test with Scaffold.

ADR-2 and ADBP-1 complex modeling:

We modeled the ADBP-1 and ADR-2 complex with AlphaFold-multimer, an AlphaFold model trained specifically for complex structure prediction (61, 62). AlphaFold-multimer significantly increases interface accuracy while maintaining high intra-chain accuracy. The AlphaFold-Multimer confidence value is defined as 0.8 · ipTM + 0.2 · pTM, where pTM (63) is a self-predicted Template Modeling score (TM-score) and ipTM is the pTM score for interface residues only. While model confidence > 0.8 is generally considered a model with a high probability of being correct, the best model of ADR-2 and ADBP-1 complex confidence value is 0.82, suggesting the model is likely accurate. To compute the interface energy for the complex, we used the pyDock scoring function based on simple but powerful electrostatics and desolvation energy terms (64). We computed the energy for all five models for the complete and mutant complexes and averaged the energy for all of them. See Supplemental Table 6.

## RESULTS

### ADR-2 nuclear localization is dependent on ADBP-1 but not on ADR-1

In *C. elegans,* most A-to-I RNA editing sites reside, as in humans, in non-coding regions, including introns, indicating that most of the editing occurs in the nucleoplasm (33, 40, 65, 66). However, direct evidence pointing to the nuclear localization of endogenous ADAR-2 is missing. Previous studies showed that *adr-2* RNA expression is high at the *C. elegans* early stages of development (40, 41, 65). Therefore, to explore the intracellular localization of ADR-2, we conducted immunostaining experiments on wild-type embryos using an antibody against ADR-2 (66). ADR-2 co-localizes with the DNA in the wild-type embryo, indicating the protein is mainly present inside nuclei (Figure 1A). Moreover, our results show that ADR-2 is adjacent to the chromosomes at all cell cycle stages (Figure 1B). Interestingly, ADR-2 is not dispersed equally along the length of the chromosomes. Previous studies showed that edited dsRNAs are enriched in the autosomal distal arm of the chromosomes, practically adjacent to repetitive sequences (33, 44, 67), with only 10.5% found in the central regions (67). In our immunostainings, ADR-2 seems to be adjacent to specific regions along the chromosomes; however, it is not only to the distal arm of the chromosomes (Figure 1B), supporting the previous study (67).

**Figure 1.**
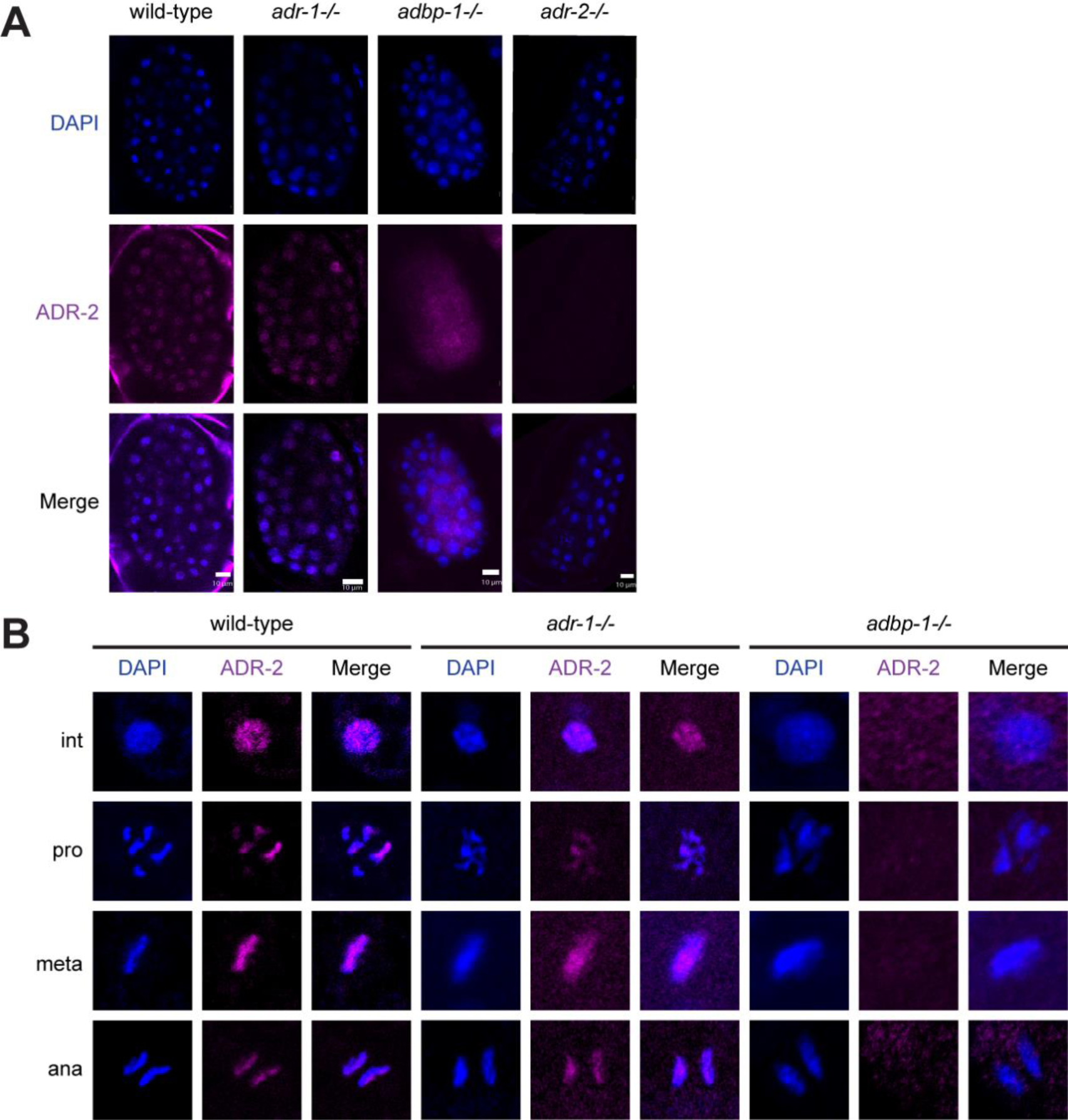
Localization of ADR-2 in embryos at different stages of mitosis. (A) Representative immunofluorescence images of DNA (DAPI in blue) and ADR-2 (magenta) in embryos from wild-type (N2), *adr-1*−/− and *adbp-1*−/− worms, including *adr-2*−/− worms as control strain. Their colocalization is shown as the overlap of both images. The surrounding background staining around the embryo in wild-type is probably a staining artifact. (B) Embryos of wild-type (N2), *adr-1*−/− and *adbp-1*−/− strains. A representative nucleus from each stage is shown: Int, interphase; Pro, prophase; Meta, metaphase; and Ana, anaphase. Scale bar, 10 µm.

Previously, it was shown that mutated *adbp-1*, truncated due to a nonsense mutation in the center of its coding region, altered the editing of four substrates and increased cytoplasmic localization of a transgenically expressed ADR-2-green fluorescent protein (GFP) driven by a hypodermal promoter (42). To test the role of ADBP-1 in the localization of the endogenous ADR-2, we examined its localization in the *adbp-1* mutant strain. Immunostaining of *adbp-1* mutant embryos with ADR-2 antibody showed that the endogenous ADR-2 is mislocalized, predominantly residing in the cytoplasm (Figure 1A, Supplemental Figure 1), in striking contrast to its nuclear location in the wild-type embryo. Comparing the average expression of ADR-2 protein in the cytoplasm and the nucleus between the *adbp-1* mutant and the wild-type shows that ADR-2 is more than two-and-a-half fold expressed in the cytoplasm of the *adbp-1* mutant than in wild-type (average (*adbp-1* mutant cytoplasm) / average (wild-type cytoplasm) = 2.5 (average *adbp-1* mutant nucleus / average wild-type nucleus), Supplemental Figure 1). The immunostaining implied that ADR-2 levels are reduced in the *adbp-1* mutant, which is consistent with our previous finding that the overall ADR-2 protein levels were slightly reduced in the *adbp-1* mutant by a western blot analysis. However, it was not completely abolished (36). In addition, ADR-2 is not localized to the chromosomes during any cell cycle stages in *adbp-1* mutant embryos (Figure 1B). To ensure that ADR-2 mislocalization is not due to a mutation in the *adr-2* gene, we sequenced the entire gene in the *adbp-1* mutant strain using several primers (Supplemental Table 1 and Supplemental Figure 2). The sequencing results showed that the *adbp-1* mutant strain has an intact *adr-2* gene with a wild-type sequence. We also verified that the *adr-2* gene is expressed (Supplemental Figure 2). To further confirm that the *adbp-1* mutation and not some background mutation causes ADR-2 mislocalization, we injected a plasmid, coding for wild-type *adbp-1* (*adbp-1p*:*gfp.adbp-1* “L” plasmid (a kind gift from Prof. Manabi Fujiwara (42))) into *adbp-1* mutant worms, generating two transgenic strains. These strains are chimeric and are not expressed in early embryos because of their extrachromosomal transgenic nature. Therefore, the effect of transgenic ADBP-1 was validated at the adult stage instead of the embryo. Nevertheless, the whole worm expression of transgenic ADBP-1 protein rescued ADR-2 nuclear localization (Supplemental Figure 3).

Because it was shown that ADR-1 regulates ADR-2 editing activity and target selection (36–38), we wondered whether ADR-1 also affects ADR-2 localization. Using immunostaining, we showed that ADR-1 and ADR-2 are mostly localized in the nuclei (Supplemental Figure 4). Immunostaining of embryos lacking *adr-1* with an ADR-2 antibody showed similar localization of ADR-2 as in wild-type worms (Figure 1A). ADR-2 localization during the cell cycle in *adr-1* mutant worms was similar to wild-type worms (Figure 1B). Therefore, we concluded that ADR-1 is not required for the nuclear localization of ADR-2.

To test if ADR-1 subcellular localization is also affected in the *adbp-1* mutant, we crossed V5-ADR-1 worms (45) to *adbp-1* mutant worms. We stained the worms with a V5 antibody. In contrast to ADR-2, in the *adbp-1* mutant, ADR-1 remains nuclear (Supplemental Figure 5); hence, ADBP-1 does not regulate ADR-1 localization.

Next, we wanted to explore ADR-2 expression during the later stages of development and in different tissues of adult worms, including germlines (Figure 2). We immunostained adult worms with the same ADR-2 antibody utilizing a technique that preserves the worm structure while allowing permeability (see Materials and methods). As a staining control, we used the MH27 antibody, which stains the apical borders of epithelial cells (Figure 2B), and the LEM-2 antibody, which stains the nuclear envelope (Supplemental Figure 6). Our results show that in wild-type adult hermaphrodites, ADR-2 is ubiquitously expressed in the intestine (Supplemental Figure 4) and gonads (Figure 2A) and not ubiquitously expressed in cells of the head, body, and tail (n=20, Figure 2B). Interestingly, in the *adbp-1* mutant adult worms, ADR-2 seems to reside more in cytoplasmic speckles of the gonads (Figure 2A), heads, and bodies of the worms (Supplemental Figure 7). Strikingly, while ADR-2 is expressed in nuclei throughout the entire oogenesis process in wild-type hermaphrodite gonads (Figure 2A), it does not appear to be expressed in sperm (Figure 2B). The lack of ADR-2 signal in the sperm supports previous research that did not identify any ADR-2 protein by sperm proteomics analysis and found only a handful of RNA molecules belonging to *adr-2* by high throughput sequencing (68). In addition to immunostaining, we also tracked RNA expression of *adr-2* and *adbp-1* in wild-type adult worms using hermaphrodite’s gonad RNA-seq data (69). We found that both genes are expressed throughout the entire oogenesis process (Supplemental Figure 8) and are not specific to a particular stage in oocyte development.

**Figure 2.**
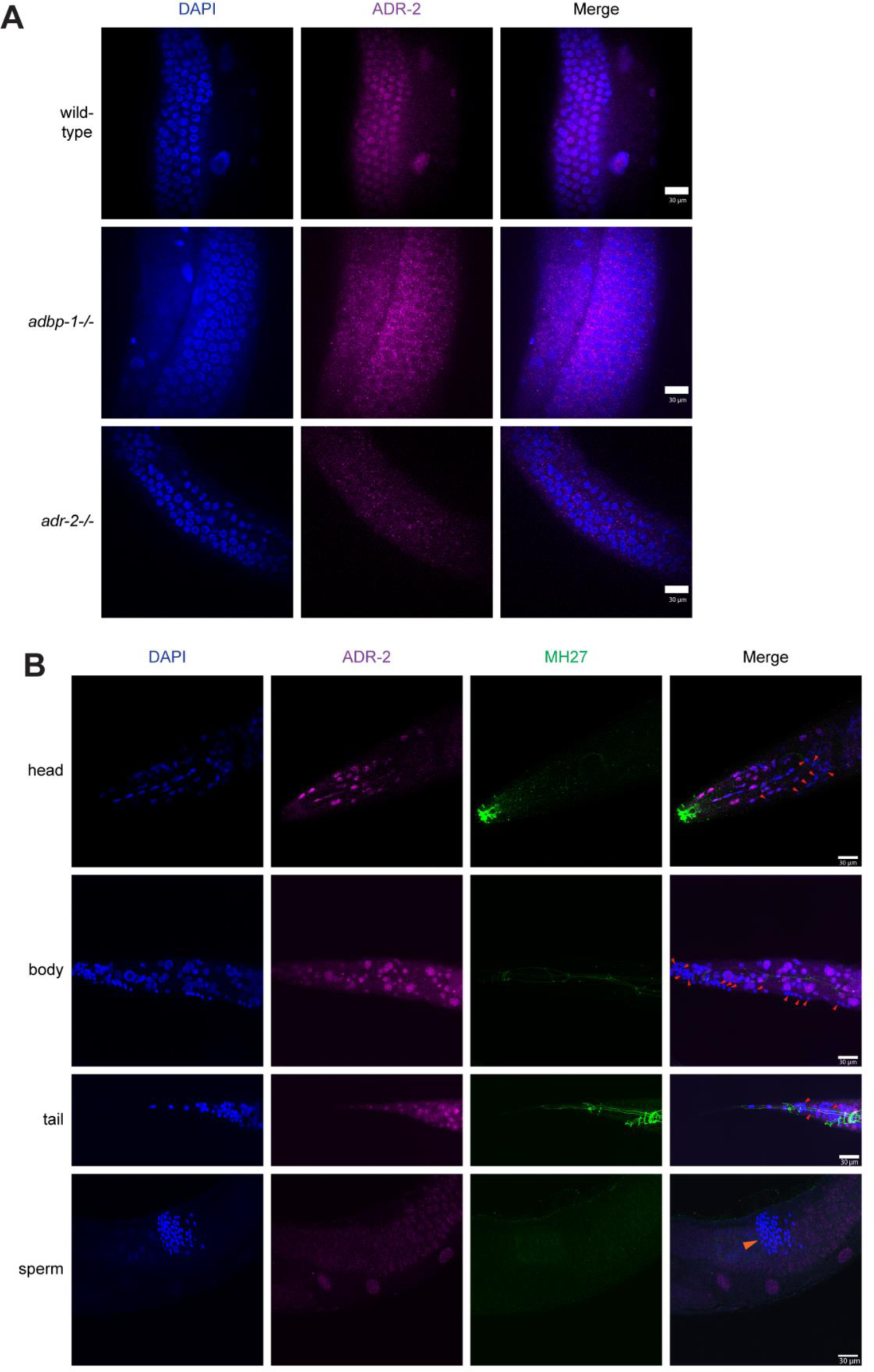
Localization of ADR-2 in the hermaphrodite adult gonads, head, body, tail, and sperm. (A) Representative immunofluorescence images of DNA (blue) and ADR-2 (magenta) from wild-type (N2) (top), *adbp-1* mutant (middle), and *adr-2*−/− control (bottom) strains. Their colocalization is shown as the overlap of both images. Scale bar, 30 µm. (B) Representative immunofluorescence images of DNA (blue), ADR-2 (magenta), and MH27 (green) from the head, body, tail, and sperm of wild-type (N2) strain. Their colocalization is shown as the overlap of the three images. Red arrows indicate where ADR-2 is absent. The orange arrow indicates the location of the sperm. Scale bar, 30 µm.

To conclude, while ADR-2 is expressed mainly in the nuclei adjacent to the chromosome through the entire cell cycle in wild-type embryos, in the *adbp-1* mutant, its expression is predominantly cytoplasmatic. ADBP-1 is required for ADR-2 nuclear localization, while ADR-1, although regulating editing performed by ADR-2, does not affect its localization. In addition, ADR-1 localization is not affected by ADBP-1. Moreover, while ADR-2 exists in the embryo and gonads, ADR-2 is not ubiquitously expressed in the worm’s later developmental stages and the sperm.

### RNA editing still occurs in the *adbp-1* mutant; however, it involves mostly coding regions

To examine if RNA editing is globally abolished in *adbp-1* mutant worms, we generated RNA-seq libraries from three biological replicas of *adbp-1* mutant worms at the embryo or L4 developmental stages. The expression data of these libraries were compared to that of libraries generated at the same developmental stages from wild-type worms (N2) and BB21 worms, which lack both *adr-1* and *adr-2* genes (*adr-1*(−/−);*adr-2*(−/−)) (40). All worms were grown under the same conditions, and most of the libraries were generated at the same time. To determine if editing occurs in the *adbp-1* mutant, we globally tested editing sites previously identified from transcriptome-wide studies (36, 40) for editing in the RNA-seq data. RNA-seq data from RNA editing mutant worms lacking *adr-1* and *adr-2* were used as a control to estimate editing false-positive sites (see Materials and Methods). We counted the number of edited sites in the wild-type and the *adbp-1* mutants (Figure 3A). As expected, in the *adbp-1* mutant, the number of edited sites is significantly reduced in both embryo and L4 developmental stages. The number of edited sites in the *adbp-1* mutant compared to wild-type worms decreased by about 70 times at the embryo stage and about 60 times at the L4 stage. However, editing is still observed (Figure 3B). We detected 188 known editing sites at the embryo stage and 124 known editing sites at the L4 stage in *adbp-1* mutant worms. Next, we wanted to examine if the distribution of editing sites in wild-type worms is different than in *adbp-1* mutant worms. In wild-type worms, almost all editing sites at both developmental stages reside in introns, as shown before (33, 40, 65, 66) (Figure 3C, D). In striking contrast, the profile of editing sites in *adbp-1* mutant worms was very different. Most editing sites in *adbp-1* mutants reside in exons in both developmental stages (Figure 3E, F). This finding aligns with our immunostaining data, which points to mainly cytoplasmic localization of ADR-2 in *adbp-1* mutant worms. As many edited genes possess several editing sites, we compared the edited genes between wild-type and *adbp-1* mutants. We found that a significant portion of the genes that undergo editing in *adbp-1* mutant worms are common to both strains (Figure 3G for the embryo, H for L4, P-value < 5.38e-47 and P-value < 1.14e-21, respectively).

**Figure 3.**
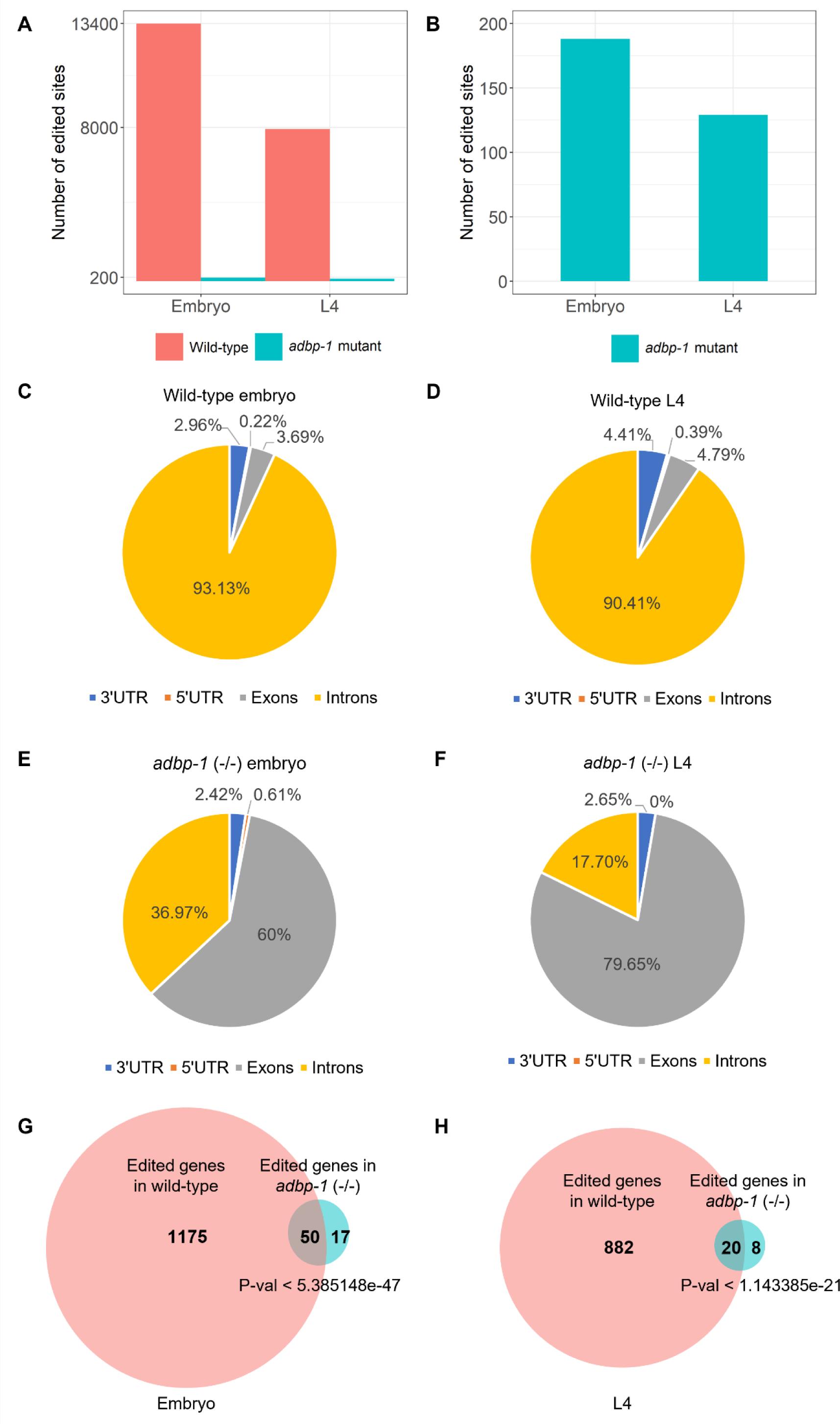
Editing sites in *adbp-1* mutant worms reside primarily in exons. (A) The bar plot represents known edited sites in the wild-type worms compared to the adbp-1 mutant worms both at the embryo stage and at the L4 larval stage. (B) The bar plot represents known edited sites in *adbp-1* mutant worms, which were not found in wild-type, at the embryo, and at the L4 larval stage. (C-F) Pie charts represent the distribution of annotated known edited sites in wild-type worms and the *adbp-1* mutant worms in the embryo and L4 stages. In the wild-type embryo worms, we detected 13398 annotated editing sites (C) and 7908 annotated editing sites at the L4 stage (D). In the *adbp-1* mutant embryo samples, we detected 188 editing sites (E), while in the *adbp-1* mutant L4 worms, we detected 129 editing sites (F). (G-H) Wild-type worms and *adbp-1* mutant worms share significant common edited genes.

After focusing on detecting editing sites found in transcriptome-wide studies (36, 40) (Figure 3), we searched if the mislocalization of ADR-2 in the cytoplasm results in *de-novo* editing sites not present in wild-type worms. To check this, we searched for new editing sites in the wild-type and *adbp-1* mutant RNA-seq using a pipeline based on the described pipelines by Goldstein et al., 2019, and Light et al., 2021 (40, 70). To be more restrictive, we considered nucleotide changes with only a Phred quality score ≥ 25 to increase the precision of nucleotide calling. We considered an edited site in the *adbp-1* mutant worms to be a *de-novo* editing site if it did not appear to be edited in wild-type worms in the current search for new editing sites or in previous works (36, 40) and appeared in at least two of the three biological replicas (see Materials and Methods and Supplemental Table 2). At the embryo stage, our pipeline detected 32 genes that had undergone editing only in *adbp-1* mutant worms. Each gene has one editing site (Figure 4A, Supplemental Table 3). At the L4 stage, 44 genes were found to be edited exclusively in *adbp-1* mutant worms, with only one editing site per gene. (Figure 4B, Supplemental Table 3). Most of the editing sites at both developmental stages reside in exons, as was shown in the analysis of known editing sites (Figure 3E, F). Interestingly, we also detected *de-novo* editing in genes normally edited in wild-type worms at the embryo and L4 stages (Figure 4A, B). At the embryo stage, only half of them reside in exons, and 43% reside in exons at the L4 stage. The *de-novo* editing sites indicate that the mislocalized ADR-2 is also active and capable of targeting transcripts but at lower levels.

**Figure 4.**
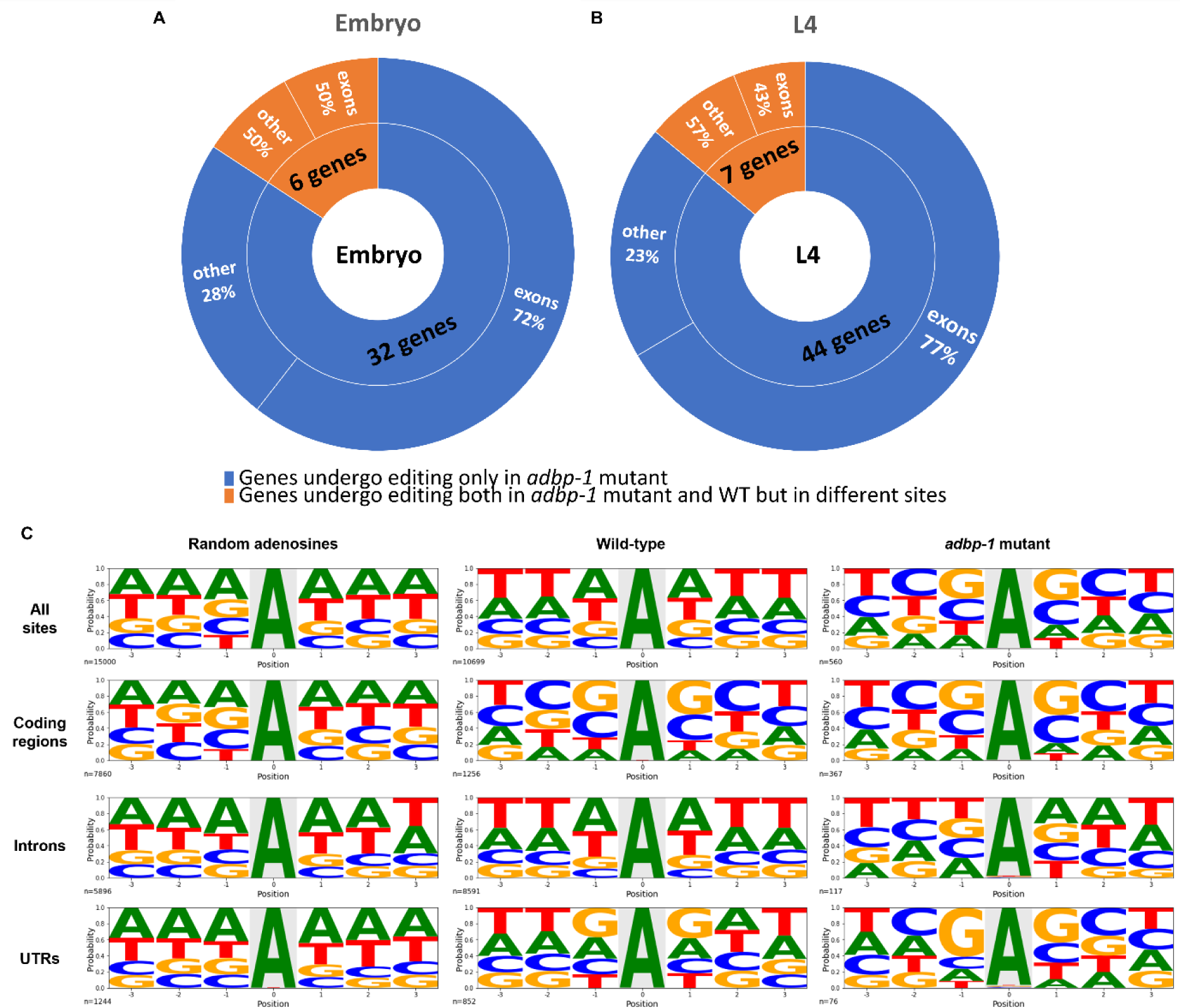
ADR-2 is enzymatically active in the cytoplasm in *adbp-1* mutant, targeting mostly exons. The pie charts describe editing sites in genes found by a pipeline searching for new editing sites in *adbp-1* mutant at the embryo (A) and L4 (B) stages. Genes found edited only in the *adbp-1* mutant and not in wild-type worms (WT) are represented by the color blue. In contrast, the orange color represents genes in *adbp-1* mutant worms that also undergo editing in wild-type worms; however, the edited sites in each gene differ between the strains. Editing sites uniquely found in *adbp-1* mutant worms tend to be in exons. (C) Distribution of nucleotides surrounding random adenosine sites and editing sites in wild-type and *adbp-1* mutant worms at coding regions, introns, UTRs, and their sums. The x-axis represents the position of the editing site (0) and its surrounding nucleotides. The y-axis represents the probability of finding each nucleotide in each position.

We wondered what defines ADR-2 targets in the *adbp-1* mutant worms. We first checked the expression level of transcripts with editing sites. We discovered that the *de-novo* edited sites in the *adbp-1* mutant belong to highly expressed protein-coding genes (Supplemental Figure 9). Therefore, it is possible that editing is not specific but guided by chance, and due to their high availability in the cytoplasm, these transcripts are being targeted by the cytoplasmic ADR-2.

Next, we wanted to make sure that editing of a site only in one of the two strains is not a result of editing levels in the second strain being below the threshold set in our pipeline (as detailed in Material and Methods) on the one hand or due to lack of expression of a transcript carrying the site in the second strain, on the other hand. For this purpose, we generated heatmaps depicting known editing sites detected in the wild-type strain and novel editing sites identified in the *adbp-1* mutant strain (Supplemental Figure 10). The heatmaps are based on calculating each nucleotide change from A-to-G and T-to-C, thus depicting all the editing events without thresholds. The heatmap discriminates between lack of editing and lack of expression. Because most editing sites in the wild-type strain are in introns, mostly remaining in nuclei, to distinguish sites existing in the nucleus and those that do not, we separated all the editing sites into 3’UTR-edited and not-3’UTR edited. Heatmaps analysis revealed that many 3’UTR-edited sites and not-3’UTR-edited that get edited in the wild-type strain remain unedited in the *adbp-1* mutant strain, albeit being expressed (Supplemental Figure 10). Similarly, most of the unique editing sites found in the *adbp-1* mutant strain are also expressed in the wild-type strain, particularly during the embryo stage, but only a small subset of them undergo editing (Supplemental Figure 10). Thus, the reason for detecting these sites only in the *adbp-1* mutant is the mislocalization of ADR-2 in *adbp-1* mutant worms and not the lack of expression in wild-type worms.

Previously, we showed that editing levels are attenuated in *adr-1* mutant worms and that ADR-1 binds RNA edited transcripts by ADR-2 directly or as part of a complex (36). Hence, ADR-1 directs ADR-2 to its editing sites. To test if ADR-1 also binds the genes edited in the *adbp-1* mutants, we divided the genes edited in the *adbp-1* mutant into known edited genes found in transcriptome-wide studies (36, 40), and uniquely (*de-novo*) edited genes found in the *adbp-1* mutant (Supplemental Figure 11). The analysis shows that a third of the known edited genes in *adbp-1* mutant worms in the embryo stage and about a third of the genes edited in the L4 stage are bound by ADR-1 (P-value = 5.52e-15 and P-value = 2.61e-09 respectively; Supplemental Figure 11A). In addition, there is no significant overlap between the newly edited genes and ADR-1-bound genes (Supplemental Figure 11B). This result suggests that ADR-1 does not guide the binding of targets by ADR-2 in the cytoplasm.

Furthermore, we aimed to understand if RNA secondary structure stability affects ADR-2 target selection in the nucleus and the cytoplasm. To check this, for each editing site found in wild-type and *adbp-1* mutant worms (known and *de-novo* editing sites), we extracted the 101 nucleotides-long sequences comprising the site and 50 surrounding nucleotides from both sides within both unspliced (nuclear) and spliced (cytoplasmatic) transcripts. As a control, we used 101 nucleotide-long sequences surrounding random adenosines. We then calculated the free energy of the secondary structure for each sequence (Supplemental Figure 12) and compared the values obtained for unspliced and spliced sequences using the Welch two-sample T-test. We found no significant differences between the secondary structures’ free energy of unspliced and spliced sequences for random adenosines and editing sites in wild-type worms. However, the free energy of unspliced sequences surrounding editing sites in *adbp-1* mutant was significantly higher than that of their spliced counterparts (Supplemental Figure 12, ΔG=−15.92 and ΔG=−17.32 correspondingly, p-value = 0.04162) and of the unspliced sequences surrounding editing sites in wild-type worms (Supplemental Figure 12, ΔG=−15.92 and ΔG=−17.67 correspondingly, p-value = 0.0343). To conclude, the secondary structure of unspliced *adbp-1* mutant edited genes is less stable than those normally edited in the wild-type nucleus and those edited in the cytoplasm when *adbp-1* is mutated.

In summary, we discovered that in the *adbp-1* mutant, editing is attenuated due to low levels of ADR-2 protein and its mislocalization. Although ADR-2 is mislocalized in the *adbp-1* mutant, it is still active, although it is not in its native environment. Two types of edited sites were detected in the *adbp-1* mutant, mostly in exons. The first type is known editing sites, present in the wild-type strain but in lower levels in the *adbp-1* mutant. The second type is *de-novo* editing sites, found only in the *adbp-1* mutant and belongs to highly-expressed coding transcripts.

### ADR-2 has a preferential site selection along the gene

Previous works showed that ADR-2 in *C. elegans* prefers adenosine or uridine as the 5′ nearest neighbor of the edited sites and adenosine as the 3’ nearest neighbor (32, 33, 71). However, the analysis was done for all sites together, regardless of their position along the gene. We wanted to understand whether neighbor nucleotides preference of editing sites residing in different regions along the gene differ and whether preferences of the *adbp-1* mutant differ from those of the wild-type worm. Hence, in our analysis, we divided edited sites found in coding genes into three groups: residing in coding regions (exons), introns, and UTRs, and performed nearest neighbor nucleotide preference analysis for each group separately. For the analysis, we used editing sites previously found in wild-type worms at all developmental stages (36, 40), editing sites we found in *adbp-1* mutant worms in this study (Figure 3B, E, F, and Figure 4A, B) and random adenosines which we randomized from different gene parts across the genome as a control. First, we analyzed all sites together, regardless of their location. As expected, the nearest neighbor nucleotide preference analysis on all sites in wild-type was similar to previous publications (32, 33), showing a preference for adenosines and uridines at the 5’ and 3’ of the editing sites. Interestingly, the nearest neighbor nucleotides of random adenosine are also adenosines (Figure 4C), probably because of the high probability of homopolymeric nucleotide runs of adenosine nucleotide in the genome (72). In contrast, we found guanosines when testing nucleotide preference at the 5’ and 3’ of the editing sites in the *adbp-1* mutant (Figure 4C). When separating editing sites according to their location along a gene, our results show that in wild-type worms, on both sides of the editing site, both in exons and UTRs, the preferred neighbor is guanosine (Figure 4C). In *adbp-1* mutant worms, the preferences in exons and UTRs are similar to those of wild-type (guanosine), while the random control shows adenosines on both sides of the selected adenosines (Figure 4C). In wild-type introns, the most common nucleotide at the 5’ and 3’ nearest neighbor positions are adenosines, as in the random control. However, in the *adbp-1* mutant, the most common nucleotide at the 5’ is uridine (shown as thymine in Figure 4C, T>G=C=A), and in the 3’, it is adenosine, as in wild-type worms (Figure 4C). However, we would like to note that the probability of a certain nucleotide might be skewed due to a very low number of editing sites residing in introns in *adbp-1* mutants. In this analysis, most editing sites (80%) in wild-type worms reside in introns. In contrast, most editing sites in *adbp-1* mutant worms (65.5%) reside in exons. Therefore, the overall editing sites motif in each strain mirrors the nucleotide preference in the region of the most abundant sites. Consequently, we conclude that the nucleotide signature preferred by ADR-2 in exons and UTRs differs from that it recognizes in exons and that the preferences are not affected by the mutation in *adbp-1*.

### 3’UTR edited genes and lncRNAs expression is altered in *adbp-1* mutant worms

Previously, we showed that 3’UTR edited genes and lncRNAs are downregulated in worms lacking *adr-2* at the embryo stage compared to wild-type worms (36, 40). This downregulation was associated with a lack of editing that allows RNAi to target substrates normally protected by editing (40). We wanted to test if the mislocalization of ADR-2 causes a similar downregulation. Using the *adbp-1* mutant RNA-seq dataset we generated, we compared the gene expression profile of *adbp-1* mutant worms to those of wild-type worms and *adr-1;adr-2* mutant worms (for this purpose, the wild-type and *adr-1;adr-2* mutant RNA-seq were re-analyzed (40)). We confirmed the downregulation of 3’UTR edited genes and lncRNA genes in *adr-1*; *adr-2* mutant worms compared to wild-type worms at the embryo stage (Figure 5A, B, P-value = 0.001 and P-value < 2.2e-16, respectively). Comparing the expression of 3’UTR edited genes and lncRNAs in *adbp-1* mutant worms to wild-type worms at the embryo stage, we saw the same trend as in *adr-1*;*adr-2* mutant worms, e.g., downregulation of 3’UTR edited genes and lncRNAs in *adbp-1* mutant worms (Figure 5C, D, P-value = 9.4e-08 and P-value < 2.2e-16, respectively). Low levels of editing in *adbp-1* mutant worms can explain this finding.

**Figure 5.**
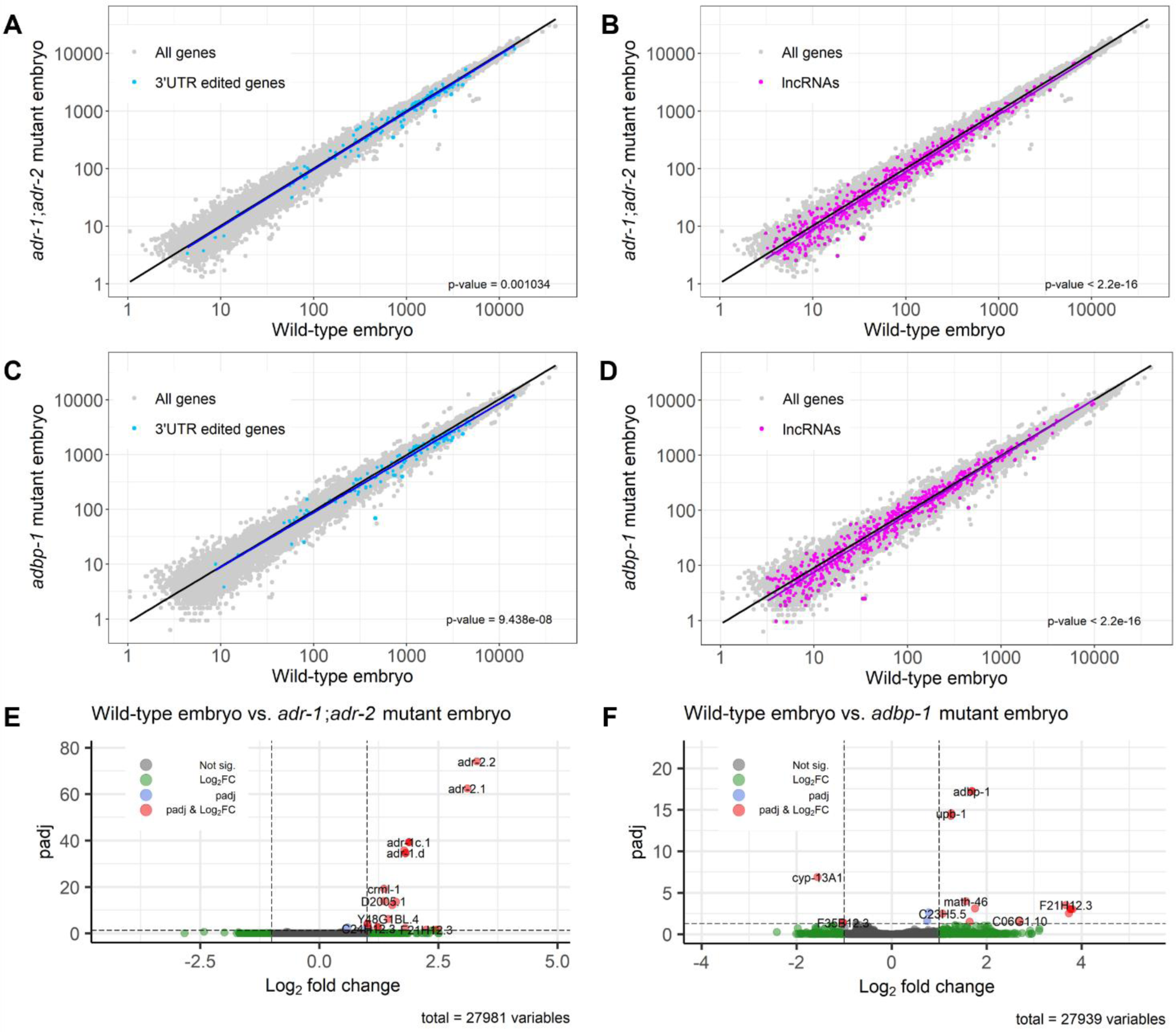
Low editing levels impair edited gene expression at the embryo stage. (A-D) The genes expressed in wild-type worms versus *adbp-1* mutant worms and the genes expressed in wild-type worms versus ADAR mutant worms, both in the embryo stage, are represented in log scale plots. Each dot represents a gene. Grey dots represent all the genes, blue dots represent edited genes at their 3’UTR, and purple dots represent lncRNAs. The black line is a regression line for all genes, the blue line is the regression line for genes edited at their 3’UTR, and the purple line is the regression line for the lncRNAs. (E-F) The volcano plots describe the log2 fold change versus −log10(P-adjusted) between the genes expressed in wild-type worms to *adr-1;adr-2* mutant worms, and wild-type worms to *adbp-1* mutant worms at the embryo. Non-significant genes are colored in grey. Differentially expressed genes, which adhere to the following criteria: |log2FoldChange| > 1 and P-adjusted < 0.05, are highlighted in red. Genes with only |log2FoldChange| > 1 are colored green, and genes with only P-adjusted < 0.05 are colored blue.

Expression analysis of 3’UTR edited genes and lncRNAs at the L4 stage was not always consistent with the embryo stage results. Previously, we reported that 3’UTR edited genes downregulated in *adr-1;adr-2* mutant worms compared to wild-type worms at the L4 stage (40). In contrast, in this analysis, 3’UTR edited genes showed slight upregulation in *adr-1;adr-2* mutant worms compared to wild-type worms. However, the P-value was close to 0.05, our significance cutoff (Supplemental Figure 13A, P-value = 0.04594). However, variations between the samples and the differences in the analysis methods might reduce the expression and significance reflected by the P-value. lncRNAs were upregulated in *adr-1;adr-2* mutant worms, compared to wild-type worms (Supplemental Figure 13B, P-value = 1.047e-10), similarly to what we showed before (40). In contrast to *adr-1*;*adr-2* mutant worms, in *adbp-1* mutant worms, 3’UTR edited genes were downregulated compared to wild-type worms (Supplemental Figure 13C, P-value = 0.0001894). At the same time, lncRNAs were not significantly changed (Supplemental Figure 13D, P-value = 0.119).

To further study ADBP-1 function, we wanted to explore the effect of lacking functional ADBP-1 on global gene expression. We wondered if the mutated *adbp-1* affects genes other than 3’UTR edited genes and lncRNAs. For this purpose, we analyzed the RNA-seq data of embryo and L4 stages, searching for differentially expressed genes. We defined a gene as differentially expressed (DE) if its expression differed more than two-fold between the two strains at the embryo stage, with a P-value after Benjamini-Hochberg correction < 0.05. Although both mutant strains showed significant downregulation of 3’UTR edited genes and lncRNAs (less than a two-fold change), we did not observe many differentially expressed genes when comparing wild-type embryos’ gene expression to *adr-1;adr-2* mutant embryos (Figure 5E, Supplemental Table 4), and when comparing wild-type embryos to *adbp-1* mutant embryos showed only a few DE genes (Figure 5F). Even when comparing *adbp-1* mutant embryos to *adr-1;adr-2* mutant embryos, the same trend of only a few DE genes was shown (Supplemental Figure 13E, Supplemental Table 4). Some DE genes found in the differential expression analysis between wild-type to *adr-1;adr-2* mutant worms were found to be DE also between wild-type to *adbp-1* mutant worms: *T11A5.8*, *R07B5.10*, *math-46*, and *F21H12.3* (Figure 5E, F). Interestingly, only *math-46* was found to be edited in wild-type worms. The three other genes, and most DE genes in the two comparisons, are neither edited in wild-type worms nor *adbp-1* mutant (Supplemental Table 4), indicating that they might be affected by upstream processes influenced by impaired editing. When we applied functional enrichment analyses of the DE of wild-type versus *adbp-1* mutant genes at the embryo stage, they showed no significant enrichment (56, 57), probably due to the low number of DE genes. When we manually examined the function of those DE genes, we saw that some are lncRNAs and some have no defined role. For a minority, their role is known, and it varies from gene to gene (73), and we could not find anything they had in common.

DE analysis of all mutant strains compared to wild-type and *adr-1*;*adr-2* mutant compared to *adbp-1* mutant worms in the L4 stage, showed much more DE genes than at the embryo stage analysis, even when we set a higher DE threshold (|log2FoldChange| > 2 and P-adjusted < 0.05, Supplemental Figure 13F-H, Supplemental Table 4). However, these variations can result from different L4 substages of the worms and not changes resulting from the difference between the strains.

To conclude, 3’UTR edited genes and lncRNAs are slightly downregulated in *adbp-1* mutant embryos compared to wild-type embryos, similar to *adr-1*;*adr-2* mutant embryos. In addition, only a few genes are significantly differentially expressed in *adbp-1* mutant embryos compared to wild-type embryos. However, some were also differentially expressed in *adr-1*;*adr-2* mutant embryos. This suggests that these genes’ expression is attenuated because of ADR-2 expression and localization changes.

### Mutated ADBP-1 has a less stable interaction with ADR-2 than the wild-type ADBP-1

In light of our results, we wanted to understand more about the interactions between ADBP-1 and ADR-2. Ohta et al. showed that ADBP-1 binds directly to ADR-2 by performing a yeast two-hybrid screen (42). To confirm this interaction and to identify additional potential ADR-2 interactors and regulators, we performed ADR-2 immunoprecipitation in wild-type worms. We subjected the precipitate to LC-MS/MS (Supplemental Table 5 and Supplemental Figure 14). Analysis of LC-MS/MS confirmed ADR-2 interactions with ADR-1 and ADBP-1 (Supplemental Table 5). Furthermore, the results point to importin proteins IMA-*3* and IMB*-3* as additional interactors (Supplemental Table 5).

ADBP-1 has no conserved domains, and *adbp-1* mutation is a nonsense mutation in the middle of the coding region (Q119STOP), shortening the protein. As we showed that the mutation can be rescued (Supplemental Figure 3), it is not a dominant negative. To better understand the mechanism of ADBP-1 and ADR-2 binding, we used AlphaFold-Multimer (61, 62) to predict their structural interaction (Figure 6A, B). AlphaFold multimer produces five high-quality models for each one of the ADBP-1-ADR-2 complexes (full-length and mutated ADBP-1). The best model for the full-length complex has a pLDDT=78.1, pTM=0.774, and ipTM=0.828 (63). Using the AlphaFold intrinsic model accuracy measure predicted TM-score (pTM) and ipTM for the interface accuracy, we got a very high model confidence measure of 0.817 (see Materials and Methods). For the truncated mutant, the best model has a pLDDT=79.8, pTM=0.812, and ipTM=0.825, and a very similar model confidence of 0.814. The confidence suggests a trustable model (see Materials and Methods). The structures modeled for the full-length and the shorter mutant complexes show very high confidence, and the relative conformation of the ADBP-1 structural is almost identical (RMSD 0.822). The prediction showed that the wild-type ADBP-1 interacts with ADR-2 at its deaminase domain (Figure 6A), physically wrapping the domain with most of its length. In truncated ADBP-1, only a small part of a protein is adjacent to ADR-2 (Figure 6B).

**Figure 6.**
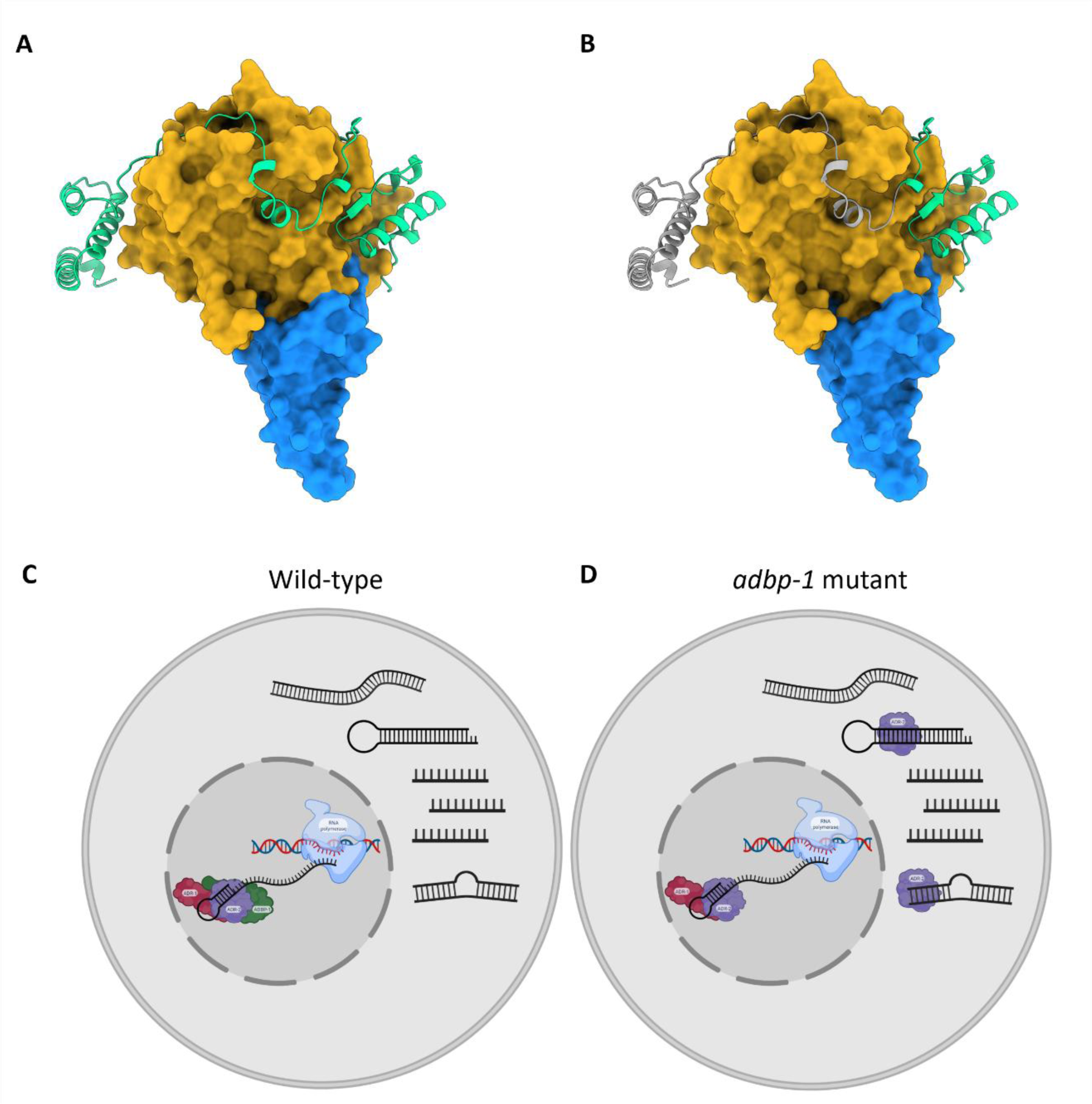
A proposed model of the interaction between ADR-2 and ADBP-1. (A) AlphaFold-Multimer prediction of ADBP-1 and ADR-2. ADBP-1 is green, the ADR-2 deaminase domain is dark yellow, and the rest of the protein is blue. (B) The mutated ADBP-1 has a missing part, noted in gray, while the remaining part is indicated in green. (C) In wild-type worms, ADBP-1 mediated ADR-2 import to the nucleus. Once ADR-2 is in the nucleus, it is adjacent to the chromosomes, where RNA editing occurs co-transcriptionally, regulated by the binding of ADR-1 to ADR-2. (D) In the *adbp-1* mutant, ADR-2 remains in the cytoplasm and edits sites randomly in exons.

To investigate further the difference between the interactions of ADR-2 with mutant and wild-type ADBP-1, we computed the total binding energy of the complex using the pyDock energy module (74) for all five models predicted. The results show that the average interface energy of the full-length ADBP-1 complex is +27 Kcal more stable than of the complex with mutant ADBP-1 (Supplemental Table 6), suggesting an important decrease in complex stability with the shorter mutant. This result suggests the mutant complex structure is less stable. Lack of stability can compromise ADBP-1’s ability to localize ADR-2 to the nucleus, which can explain our experimental data.

Taking together all of our findings, we propose the following model. In wild-type worms, full-length ADBP-1 interacts with ADR-2 by wrapping itself around its deaminase domain. This interaction mediates ADR-2 import to the nucleus. In the nucleus, ADR-2 is directed by ADR-1 to the specific editing sites in dsRNA molecules. (Figure 6C, D). The proximity of ADR-2 to chromosomes enables cotranscriptional editing and nascent RNA binding. Truncation mutation in ADBP-1 affects the stability of its complex with ADR-2, affecting ADBP-1 functionality. Without functional ADBP-1 to guide it into the nucleus, ADR-2 remains mainly in the cytoplasm, where it edits spliced transcripts. Hence, most editing happens in exons. Editing in cytoplasm is probably not guided by ADR-1. It occurs sporadically in highly expressed transcripts possessing dsRNA regions. Interestingly, the nucleotide signature surrounding editing sites differs for sites in the introns, exons, and UTRs (Figure 4C).

## DISCUSSION

In this work, we sought to examine the intracellular and tissue-specific localization of ADR-2 in *C. elegans* and the effect the localization might have on its function. Although the general assumption is that editing occurs mainly in the nucleus (42, 65), surprisingly, the exact location of the endogenous ADR-2 protein in *C. elegans* has not been directly shown. Using a specific antibody against ADR-2, we showed that in wild-type worms, ADR-2 resides in nuclei and is adjacent to the chromosomes at all cell cycle stages.

We use *adbp-1* and *adr-1* mutant worms to show that ADBP-1, not ADR-1, facilitates endogenous ADR-2 localization to the nucleus. In the absence of ADBP-1, ADR-2 appears in the cytoplasm. We also show that ADBP-1 does not affect ADR-1 localization, which is mainly nuclear. In addition, our bioinformatical analysis revealed that although the editing level decreased in the absence of ADBP-1, ADR-2 is still enzymatically active. Worms lacking *adbp-1* still exhibit non-negligible editing, mainly in exons, probably due to their cytoplasmic localization. Many genes undergo editing in both wild-type and *adbp-1* mutant worms.

Moreover, the mislocalization of ADR-2 leads to *de-novo* editing sites that do not exist in wild-type worms. *De-novo* editing appeared in highly expressed genes that were not found to be bound by ADR-1, indicating sporadic editing of ADR-2 in the cytoplasm. Our bioinformatical analysis showed that ADR-2 preferences for nucleotides surrounding targeted adenosines in exons, introns, and UTRs differ.

Looking at RNA expression levels in ADBP-1 mutant embryos, we noticed the downregulation of genes that undergo editing in their 3’UTR normally and lncRNAs. Similar downregulation, previously observed by us in ADAR mutant worms (40), is attributed to the sensitivity of unedited transcripts to RNAi. ADR-2 is highly expressed in all embryo cells, as shown before (40, 65). In contrast, it is not ubiquitously expressed in the somatic tissues of adults. In wild-type adult hermaphrodites, ADR-2 is expressed mainly in the gonad (Figure 2A), not in sperm (Figure 2B).

### ADR-2 nuclear localization proximity to the chromosomes suggests cotranscriptional editing

In the embryo, we showed that ADR-2 is expressed in the nuclei of most cells (Figure 1A), which is consistent with research showing that the expression of *adr-2* mRNA is highest at the early developmental stages (40, 65).

Not only does ADR-2 reside in the nuclei in embryos, but it also seems to localize near the chromosomes (Figure 1B). This proximity could allude to the importance of ADR-2 being close to the DNA so that the editing process can happen co-transcriptionally when transcription begins or ADR-2 binding to nascent transcripts. In addition, most of the editing sites in *C. elegans* being in introns (65), also suggesting cotranscriptional editing. This finding aligns with other research showing cotranscriptional RNA editing in humans (75, 76) and *Drosophila* (77). The study in humans revealed that A-to-I RNA editing events occur in nascent RNA associated with chromatin before polyadenylation (75, 76).

In addition to ADR-2 chromosome proximity, we observed that ADR-2 is not distributed evenly along chromosomes but is localized in specific regions (Figure 1B). Previous studies in *C. elegans* showed that autosomal chromosomes distal arms are enriched with dsRNA, practically in repetitive sequences (33, 44, 67). Although it is difficult to differentiate the exact localization of ADR-2 along the chromosomes in our results, ADR-2 may also be localized at the autosome distal arm and co-transcriptionally edits its targets. The localization of ADR-2 in the absence of ADR-1 is identical to that of the wild-type. This points to two independent stages in localizing ADR-2; in the first, ADR-2 is brought to chromosomes (and possibly to the particular chromosome areas) by regulator/s other than ADR-1, and, at the second stage, ADR-2 is targeted by ADR-1 to its specific editing substrates. This suggests that more regulators of this process remain to be discovered.

### ADBP-1 regulates the nuclear localization of ADR-2 and not of ADR-1

In human, both ADAR1 isoforms, 110-kDa, and 150-kDa protein, can shuttle between the nucleus and cytoplasm. However, ADAR1 p150 has a strong nuclear export signal, overlapping its third dsRBD, which leads to its accumulation in the cytoplasm (20, 21). In contrast, human ADAR2 is mainly localized to the nucleolus (24). Human ADAR2 localization is regulated by binding to pin1 (78). In *C. elegans*, ADBP-1 was shown to regulate the localization of the transgenic ADR-2 (42). In contrast to using a transgenic ADR-2, we aimed to understand how ADBP-1 affects endogenous ADR-2. We confirmed the finding that ADBP-1 regulates the nuclear localization of *C. elegans* ADR-2 (Figures 1, 2). ADR-1 was shown to regulate editing by ADR-2, probably by directing ADR-2 to the editing sites (36–38). Our results suggest that ADR-1 regulates editing but does not affect ADR-2 localization. It is still unclear whether ADBP-1 affects ADR-2 localization directly or indirectly. To understand more about *C. elegans* ADR-2 shuttling, we tried to predict the existence of nuclear localization signal (NLS) in ADR-2 using NLS prediction tools (79–82); however, we could not find any NLS in ADR-2. Though, these tools are limited because they mainly cover classical NLSs, not accounting for non-classical ones, as in human ADAR1 (19).

Interestingly, we could not detect an NLS in ADBP-1 as well. ADBP-1 may have a non-classical NLS or serve as an adaptor protein, mediating the active import of ADR-2 to the nucleus (Figure 6C, D). Alternatively, a lack of NLS can suggest that ADR-2 and ADBP-1 may also have a cytoplasmic activity, which can be expressed only in specific cells or tissues. In such a case, the localization of ADR-2 probably regulates its function.

Although we could not detect an NLS in ADR-2 and ADBP-1, we found that ADR-2 interacts with the importins IMA-3 and IMB-3 (Supplemental Table 5). These importins may facilitate the transport of ADR-2 to the nucleus. In light of the results showing that in the *adbp-1* mutant, ADR-2 resides in the cytoplasm, it is possible that IMA-3 and IMB-3 bind ADR-2 only when it is bound to ADBP-1. We showed that the binding of ADR-2 with wild-type ADBP-1 is more stable than with a mutated ADBP-1 (Supplemental Table 6). Hence, it is possible that in case of unstable binding of ADR-2 with mutated ADBP-1, IMA-3 and IMB-3 cannot bind the ADR-2-ADBP-1 complex, causing ADR-2 to remain in the cytoplasm.

Moreover, it is known that small water-soluble molecules weighing less than ∼60 kDa can diffuse into the nucleus (83). As ADR-2 has a molecular weight of 55 kDa (Supplemental Table 5), it is possible that a NLS is not required for ADR-2 as it can passively diffuse through nuclear pore complexes (NPCs). In such case, in the absence of functional ADBP-1, ADR-2 can be present in a nucleus but at lower levels. In the nucleus, it can bind to ADR-1, which is not affected by ADBP-1 (Supplemental Figure 5), and target genes. Indeed, we found a fraction of intron-residing editing sites to undergo editing in ADBP-1 mutant worms (Figure 3E-F and Figure 4A-B). In addition, when we analyzed if *adbp-1* mutant worms have editing in genes previously identified from transcriptome-wide studies (36, 40) (Figure 3A, B), we found that although the small number of edited sites, a significant fraction of these edited genes are bound by ADR-1 (Supplemental Figure 11A).

We previously showed that ADR-1 binds dsRNA at editing sites (36). In addition, ADR-2 has a low affinity to dsRNA, which increases upon its binding to ADR-1 (38). Thus, after ADBP-1 brings ADR-2 to the nucleus, the binding of ADR-2 to ADR-1 brings ADR-2 to its proper RNA targets. The cytoplasmic location of ADR-2 leads to different editing patterns for several reasons. High cytoplasmic levels of ADR-2 result in editing in the cytoplasm, mainly in exons, including novel sites. On the other hand, low ADR-2 levels in the nucleus result in a lack of editing in introns (See our model Figure 6C, D). In addition, in the cytoplasm, the lack of introns in the transcripts decreases dsRNA structures, which are ADR-2 substrates. Hence, editing levels drop and mainly occur in exons. Despite the fact that we found a significant fraction of *adbp-1* mutant edited genes are bound by ADR-1 (Supplemental Figure 11A), it is possible that the ADR-2 reduced levels in the nucleus due to the mutation in *adbp-1* (Supplemental Figure 1 and Supplemental Figure 3B) could also change the amount and nature of ADR-1-bound transcripts. A decreasion of transcripts bound by ADR-1 can cause ADR-2 not to target them.

The *adbp-1* mutation downregulates the expression of 3’UTR-edited genes and lncRNAs in embryos (Figure 5A, B). This can be explained by their failure to undergo editing in the nucleus, like introns. Because of long unedited dsRNA stretches, 3’UTR edited genes and lncRNAs are successfully targeted by RNAi machinery, leading to the observed downregulation of their levels, similar to what happens in ADARs mutants. In addition, ADR-2 and ADBP-1 may have additional roles in the cell, which leads to downstream regulation of these genes and the DE genes we found (Figure 5E, F and Supplemental Figure 13E-H).

### ADR-2 expression pattern in embryo and adult worm is different

In contrast to the embryo, ADR-2 expression is not evident in all cells of somatic tissues of the worm’s head, body, and tail (Figure 2B). In humans, ADAR proteins, mainly ADAR2, have essential functions in the nervous system (9–12). In *C. elegans,* neuronal genes were found to undergo editing (40, 66), and one of the most prominent phenotypes of worms lacking *adr-2* is chemotaxis defects (34, 66), which may be explained by the expression of ADR-2 seen in neuronal cells (34, 66),

The ubiquitous presence of ADR-2 in the oocytes indicates that RNA editing is needed for the entire developmental process, which starts with oocyte development and maturation and continues after fertilization into embryonic development. This could be consistent with other studies showing that editing levels are highest during *C. elegans* earlier stages of development (embryo and L1) (40, 44, 65). In a striking difference to a strong expression of ADR-2 in the gonad (Figure 2A, B), we could not detect ADR-2 in sperm. The existence of ADR-2 only in certain tissues and cells indicates a control that occurs not only at the intracellular level but also at the tissue level.

### What defines ADR-2 targets?

The cytoplasmic editing in exons in *adbp-1* mutant belongs to highly expressed genes (Supplemental Figure 9). Therefore, we assume that because ADR-1 resides mostly in the nucleus, ADR-2 targets these genes because of their abundance, increasing the chances of ADR-2 encountering and targeting them.

When we tried to characterize the stability of the dsRNAs targeted by ADR-2 (Supplemental Figure 12), we found that genes targeted in the wild-type have no significant difference in their stability, whether spliced or not. However, because most wild-type editing sites reside in introns (Figure 3C, D), we can conclude that ADR-2 targets unspliced sequences with ADR-1 guidance. As we suggested before, editing may happen co-transcriptionally. Interestingly, when we checked genes edited in the *adbp-1* mutant, we found that their unspliced form is significantly less stable than wild-type edited unspliced genes (Supplemental Figure 12). Hence, it is possible that when ADR-2 is mainly in the cytoplasm, it encounters high abundance dsRNAs that are more stable than their unspliced form in the nucleus and, therefore, are not normally edited.

While the previous works analyzed the nucleotides surrounding the targeted adenosine and looked at the overall editing sites, we focused on nucleotides surrounding edited adenosines in UTRs, coding exons, and introns separately (Figure 4C). By looking at the nucleotides surrounding the overall edited adenosine that appears at all genes’ parts together, our results align with previous works done in *C. elegans* (36, 40). Surprisingly, when we focused on the different gene parts, we found different nucleotides surrounding the editing site in each gene part. The specific editing motif of each gene part was very similar in the wild-type and the *adbp-1* mutant. This suggests that ADR-2 cellular localization does not affect the motif. Most editing sites we analyzed in the *adbp-1* mutant are not novel and were found in transcriptome-wide studies in different developmental stages in wild-type worms (Figure 3). Hence, we assume they probably edited in the mutant when ADR-2 exists in the nucleus with the ADR-1. We do not know what makes ADR-2 prefer specific nucleotides along the gene or what causes ADR-1 to direct ADR-2 to specific adenosines. Still, more unknown factors may facilitate this process and guide ADR-2 to the different motifs.

## AVAILABILITY

The sequence data from this study have been submitted to the NCBI Gene Expression Omnibus (GEO; http://www.ncbi.nlm. nih.gov/geo/) under accession number GSE230883.

## ACKNOWLEDGEMENTS

We thank Dr. Alla Fishman for the critical reading of the manuscript. We want to thank Brenda Bass for providing strains and Prof. Benjamin Podbilewicz for the immunostaining antibodies. We thank Prof. Manabi Fujiwara for providing the *adbp-1 gfp* “L” plasmid. In addition, we thank Dr. Menachem Katz for helping with worm injections and Dr. Clari Valensi for helping with the immunostaining experiments. Some strains were provided by the CGC, which is funded by NIH Office of Research Infrastructure Programs (P40 OD010440). Figure 6C, D and the graphical abstract were created with BioRender.com.

## REFERENCES

1. Gott, J.M. and Emeson, R.B. (2000) Functions and Mechanisms of RNA Editing. Annu. Rev. Genet. 34, 499–531.

2. Erdmann, E.A., Mahapatra, A., Mukherjee, P., Yang, B. and Hundley, H.A. (2021) To protect and modify double-stranded RNA - the critical roles of ADARs in development, immunity and oncogenesis. Crit. Rev. Biochem. Mol. Biol., 56, 54–87.

3. Walkley, C.R. and Li, J.B. (2017) Rewriting the transcriptome: adenosine-to-inosine RNA editing by ADARs. Genome Biol., 18, 205.

4. Levanon, Y.E. and Eisenberg, E. (2018) A-to-I RNA editing — immune protector and transcriptome diversifier. Nat. Rev. Genet., 19, 473–490.

5. Bass, B.L. (2002) RNA Editing by Adenosine Deaminases That Act on RNA. Annu. Rev. Biochem., 71, 817–846.

6. Quin, J., Sedmík, J., Vukić, D., Khan, A., Keegan, L.P. and O’Connell, M.A. (2021) ADAR RNA Modifications, the Epitranscriptome and Innate Immunity. Trends Biochem. Sci., 46, 758–771.

7. Higuchi, M., Single, F.N., Kohler, M., Sommer, B., Sprengel, R. and Seeburg, P.H. (1993) RNA Editing of AMPA Receptor Subunit GluR-9: A Base-Paired Intron-Exon Structure Determines Position and Efficiency. Cell, 75, 1361–1370.

8. Paz, N., Levanon, E.Y., Amariglio, N., Heimberger, A.B., Ram, Z., Constantini, S., Barbash, Z.S., Adamsky, K., Safran, M., Hirschberg, A., et al. (2007) Altered adenosine-to-inosine RNA editing in human cancer. Genome Biol., 17(11):1586–95.

9. Kawahara, Y., Ito, K., Sun, H., Aizawa, H., Kanazawa, I. and Kwak, S. (2004) RNA editing and death of motor neurons. Nature, 427, 6977:801.

10. Jain, M., Jantsch, M.F. and Licht, K. (2019) The Editor’s I on Disease Development. Trends Genet., 35, 903–913.

11. Song, B., Shiromoto, Y., Minakuchi, M. and Nishikura, K. (2022) The role of RNA editing enzyme ADAR1 in human disease. Wiley Interdiscip. Rev. RNA, 13, 1–35.

12. Yang, Y., Okada, S. and Sakurai, M. (2021) Adenosine-to-inosine RNA editing in neurological development and disease. RNA Biol., 18, 999–1013.

13. Baker, A.R. and Slack, F.J. (2022) ADAR1 and its implications in cancer development and treatment. Trends Genet., 38, 821–830.

14. Shiromoto, Y., Sakurai, M., Minakuchi, M., Ariyoshi, K. and Nishikura, K. (2021) ADAR1 RNA editing enzyme regulates R-loop formation and genome stability at telomeres in cancer cells. Nat. Commun., 12(1):1654.

15. Steele, E.J. and Lindley, R.A. (2017) ADAR deaminase A-to-I editing of DNA and RNA moieties of RNA:DNA hybrids has implications for the mechanism of Ig somatic hypermutation. DNA Repair (Amst)., 55, 1–6.

16. Zheng, Y., Lorenzo, C. and Beal, P.A. (2017) DNA editing in DNA/RNA hybrids by adenosine deaminases that act on RNA. Nucleic Acids Res., 45, 3369–3377.

17. Jimeno, S., Prados-Carvajal, R., Fernández-Ávila, M.J., Silva, S., Silvestris, D.A., Endara-Coll, M., Rodríguez-Real, G., Domingo-Prim, J., Mejías-Navarro, F., Romero-Franco, A., et al. (2021) ADAR-mediated RNA editing of DNA:RNA hybrids is required for DNA double strand break repair. Nat. Commun., 12(1):5512.

18. Patterson, J.B. and Samuel, C.E. (1995) Expression and regulation by interferon of a double-stranded-RNA-specific adenosine deaminase from human cells: evidence for two forms of the deaminase. Mol. Cell. Biol., 15, 5376–5388.

19. Barraud, P., Banerjee, S., Mohamed, W.I., Jantsch, M.F. and Allain, F.H. (2014) A bimodular nuclear localization signal assembled via an extended double-stranded RNA-binding domain acts as an RNA-sensing signal for transportin 1. PNAS., 111(18):E1852–61.

20. Fritz, J., Strehblow, A., Taschner, A., Schopoff, S., Pasierbek, P. and Jantsch, M.F. (2009) RNA-Regulated Interaction of Transportin-1 and Exportin-5 with the Double-Stranded RNA-Binding Domain Regulates Nucleocytoplasmic Shuttling of ADAR1. Mol. Cell. Biol., 29, 1487–1497.

21. Poulsen, H., Nilsson, J., Damgaard, C.K., Egebjerg, J. and Kjems, J. (2001) CRM1 Mediates the Export of ADAR1 through a Nuclear Export Signal within the Z-DNA Binding Domain. Mol. Cell. Biol., 21, 7862–7871.

22. Patterson, J.B. and Samuel, C.E. (1995) Expression and regulation by interferon of a double-stranded-RNA-specific adenosine deaminase from human cells: evidence for two forms of the deaminase. Mol. Cell. Biol., 15, 5376–5388.

23. Kleinova, R., Leuchtenberger, A.F., Giudice, C. Lo, Tanzer, A., Derdak, S., Picardi, E. and Jantsch, M.F. (2023) The ADAR1 editome reveals drivers of editing-specificity for ADAR1-isoforms. Nucleic Acids Res. 51(9):4191–4207.

24. Desterro, J.M.P., Keegan, L.P., Lafarga, M., Berciano, M.T., O’Connell, M. and Carmo-Fonseca, M. (2003) Dynamic association of RNA-editing enzymes with the nucleolus. J. Cell Sci., 116, 1805–1818.

25. Higuchi, M., Maas, S., Single, F.N., Hartner, J., Rozov, A., Burnashev, N., Feldmeyer, D., Sprengel, R. and Seeburg, P.H. (2000) Point mutation in an AMPA receptor gene rescues lethality in mice deficient in the RNA-editing enzyme ADAR2. Nature, 406, 1998–2001.

26. Roth, S.H., Levanon, E.Y. and Eisenberg, E. (2019) Genome-wide quantification of ADAR adenosine-to-inosine RNA editing activity. Nat. Methods, 16(11):1131–1138.

27. Eggington, J.M., Greene, T. and Bass, B.L. (2011) Predicting sites of ADAR editing in double-stranded RNA. Nat Commun, 2, 319.

28. Lehmann, K.A. and Bass, B.L. (2000) Double-stranded RNA adenosine deaminases ADAR1 and ADAR2 have overlapping specificities. Biochemistry, 39, 12875–12884.

29. Polson, A.G. and Bass, B.L. (1994) Preferential selection of adenosines for modification by double-stranded RNA adenosine deaminase. EMBO J., 13, 5701–5711.

30. Riedmann, E.M., Schopoff, S., Hartner, J.C. and Jantsch, M.F. (2008) Specificity of ADAR-mediated RNA editing in newly identified targets. Rna, 14, 1110–1118.

31. Morse, D.P. and Bass, B.L. (1999) Long RNA hairpins that contain inosine are present in Caenorhabditis elegans poly(A)+ RNA. Proc Natl Acad Sci U S A, 96, 6048–6053.

32. Washburn, M.C., Kakaradov, B., Sundararaman, B., Wheeler, E., Hoon, S., Yeo, G.W. and Hundley, H.A. (2014) The dsRBP and inactive editor, ADR-1, utilizes dsRNA binding to regulate A-to-I RNA editing across the C. elegans transcriptome. Cell Rep., 6, 599–607.

33. Whipple, J.M., Youssef, O.A., Aruscavage, P.J., Nix, D.A., Hong, C., Johnson, W.E. and Bass, B.L. (2015) Genome-wide profiling of the C. elegans dsRNAome. Rna, 21, 786–800.

34. Tonkin, L.A., Saccomanno, L., Morse, D.P., Brodigan, T., Krause, M. and Bass, B.L. (2002) RNA editing by ADARs is important for normal behavior in Caenorhabditis elegans. EMBO J, 21, 6025–6035.

35. Arribere, J.A., Kuroyanagi, H. and Hundley, H.A. (2020) mRNA editing, processing and quality control in caenorhabditis elegans. Genetics, 215, 531–568.

36. Ganem, N.S., Ben-Asher, N., Manning, A.C., Deffit, S.N., Washburn, M.C., Wheeler, E.C., Yeo, G.W., Zgayer, O.B.N., Mantsur, E., Hundley, H.A., et al. (2019) Disruption in A-to-I Editing Levels Affects C. elegans Development More Than a Complete Lack of Editing. Cell Rep., 27, 1244–1253.e4.

37. Washburn, M.C., Kakaradov, B., Sundararaman, B., Wheeler, E., Hoon, S., Yeo, G.W. and Hundley, H.A. (2014) Report The dsRBP and Inactive Editor ADR-1 Utilizes dsRNA Binding to Regulate A-to-I RNA Editing across the C.elegans Transcriptome. CellReports, 6, 599–607.

38. Rajendren, S., Manning, A.C., Al-Awadi, H., Yamada, K., Takagi, Y. and Hundley, H.A. (2018) A protein-protein interaction underlies the molecular basis for substrate recognition by an adenosine-to-inosine RNA-editing enzyme. Nucleic Acids Res., 46(18):9647–9659.

39. Rajendren, S., Dhakal, A., Vadlamani, P., Townsend, J., Deffit, S.N. and Hundley, H.A. (2021) Profiling neural editomes reveals a molecular mechanism to regulate RNA editing during development. Genome Res., 31, 27–39.

40. Goldstein, B., Agranat-Tamir, L., Light, D., Zgayer, O.B.N., Fishman, A. and Lamm, A.T. (2017) A-to-I RNA editing promotes developmental stage-specific gene and lncRNA expression. Genome Res., 27, 462–470.

41. Hundley, H.A., Krauchuk, A.A. and Bass, B.L. (2008) C. elegans and H. sapiens mRNAs with edited 3′ UTRs are present on polysomes. Rna, 14, 2050–2060.

42. Ohta, H., Fujiwara, M., Ohshima, Y. and Ishihara, T. (2008) ADBP-1 Regulates an ADAR RNA-Editing Enzyme to Antagonize RNA-Interference-Mediated Gene Silencing in Caenorhabditis elegans. Genetics, 180(**2**), 785–796.

43. Brenner, S. (1974) The Genetics of Caenorhabditis Elegans. Genetics, 4, 71–94.

44. Reich, D.P., Tyc, K.M. and Bass, B.L. (2018) C. elegans ADARs antagonize silencing of cellular dsRNAs by the antiviral RNAi pathway. Genes Dev., 32, 271–282.

45. Dhakal, A., Salim, C., Skelly, M., Amichan, Y., Lamm, A.T. and Hundley, H.A. (2024) ADARs regulate cuticle collagen expression and promote survival to pathogen infection. BMC Biol., 22, 1–17.

46. Williams, B.D., Schrank, B., Huynh, C., Shownkeen, R. and Waterston, R.H. (1992) A genetic mapping system in Caenorhabditis elegans based on polymorphic sequence-tagged sites. Genetics, 131, 609–624.

47. Andrews, Simon. FastQC: a quality control tool for high throughput sequence data. 2010. (2017): W29–33.

48. Langmead, B., Trapnell, C., Pop, M. and Salzberg, S.L. (2009) Ultrafast and memory-efficient alignment of short DNA sequences to the human genome. Genome Biol., 10.

49. Li, H., Handsaker, B., Wysoker, A., Fennell, T., Ruan, J., Homer, N., Marth, G., Abecasis, G. and Durbin, R. (2009) The Sequence Alignment/Map format and SAMtools. Bioinformatics, 25, 2078–2079.

50. Harris, T.W., Antoshechkin, I., Bieri, T., Blasiar, D., Chan, J., Chen, W.J., De La Cruz, N., Davis, P., Duesbury, M., Fang, R., et al. (2010) WormBase: a comprehensive resource for nematode research. Nucleic Acids Res., 38, D463–D467.

51. Howe, K.L., Bolt, B.J., Cain, S., Chan, J., Chen, W.J., Davis, P., Done, J., Down, T., Gao, S., Grove, C., et al. (2016) WormBase 2016: Expanding to enable helminth genomic research. Nucleic Acids Res., 44, D774–D780.

52. Howe, K.L., Bolt, B.J., Shafie, M., Kersey, P. and Berriman, M. (2017) WormBase ParaSite − a comprehensive resource for helminth genomics. Mol. Biochem. Parasitol., 215, 2–10.

53. Tareen, A. and Kinney, J.B. (2020) Logomaker: Beautiful sequence logos in Python. Bioinformatics, 36, 2272–2274.

54. Love, M.I., Huber, W. and Anders, S. (2014) Moderated estimation of fold change and dispersion for RNA-seq data with DESeq2. Genome Biol., 15, 1–21.

55. Blighe K, Rana S, L.M. (2022) EnhancedVolcano: Publication-ready volcano plots with enhanced colouring and labeling.

56. Angeles-Albores, D., Raymond, R.Y., Chan, J. and Sternberg, P.W. (2016) Tissue enrichment analysis for C. elegans genomics. BMC Bioinformatics, 17, 1–10.

57. Angeles-Albores, D., Lee, R., Chan, J. and Sternberg, P. (2018) Two new functions in the WormBase Enrichment Suite. microPublication Biol., 10.17912/W25Q2N.

58. Margalit, A., Segura-Totten, M., Gruenbaum, Y. and Wilson, K.L. (2005) Barrier-to-autointegration factor is required to segregate and enclose chromosomes within the nuclear envelope and assemble the nuclear lamina. Proc. Natl. Acad. Sci. U. S. A., 102, 3290–3295.

59. Finney, M. and Ruvkun, G. (1990) The unc-86 gene product couples cell lineage and cell identity in C. elegans. Cell, 63, 895–905.

60. Schneider, C.A., Rasband, W.S. and Eliceiri, K.W. (2012) NIH Image to ImageJ: 25 years of image analysis. Nat. Methods, 9, 671–675.

61. Jumper, J., Evans, R., Pritzel, A., Green, T., Figurnov, M., Ronneberger, O., Tunyasuvunakool, K., Bates, R., Žídek, A., Potapenko, A., et al. (2021) Highly accurate protein structure prediction with AlphaFold. Nature, 596, 583–589.

62. Evans, R., O’Neill, M., Pritzel, A., Antropova, N., Senior, A., Green, T., Žídek, A., Bates, R., Blackwell, S., Yim, J., et al. (2022) Protein complex prediction with AlphaFold-Multimer. bioRxiv, 10.1101/2021.10.04.463034.

63. Zhang, Y. and Skolnick, J. (2005) TM-align: A protein structure alignment algorithm based on the TM-score. Nucleic Acids Res., 33, 2302–2309.

64. Grosdidier, S., Pons, C., Solernou, A. and Fernández-Recio, J. (2007) Prediction and scoring of docking poses with pyDock. Proteins, 69, 852–858.

65. Zhao, H.Q., Zhang, P., Gao, H., He, X., Dou, Y., Huang, A.Y., Liu, X.M., Ye, A.Y., Dong, M.Q. and Wei, L. (2015) Profiling the RNA editomes of wild-type C. elegans and ADAR mutants. Genome Res, 25, 66–75.

66. Deffit, S.N., Yee, B.A., Manning, A.C., Rajendren, S., Vadlamani, P., Wheeler, E.C., Domissy, A., Washburn, M.C., Yeo, G.W. and Hundley, H.A. (2017) The C. elegans neural editome reveals an ADAR target mRNA required for proper chemotaxis. Elife, 6, e28625.

67. Reich, D.P. and Bass, B.L. (2018) Inverted repeat structures are associated with essential and highly expressed genes on C. Elegans autosome distal arms. Rna, 24, 1634–1646.

68. Ma, X., Zhu, Y., Li, C., Xue, P., Zhao, Y., Chen, S., Yang, F. and Miao, L. (2014) Characterisation of Caenorhabditis elegans sperm transcriptome and proteome. BMC Genomics, 15, 1–13.

69. Yonatan B. Tzur, Eitan Winter, Jinmin Gao, Tamar Hashimshony, Itai Yanai, and M.P.C. (2018) Spatiotemporal Gene Expression Analysis of the Caenorhabditis elegans Germline Uncovers a Syncytial Expression Switch. Genetics, 210, 587–605.

70. Light, D., Haas, R., Yazbak, M., Elfand, T., Blau, T. and Lamm, A.T. (2021) RESIC: A Tool for Comprehensive Adenosine to Inosine RNA Editing Site Identification and Classification. Front. Genet., 12, 686851.

71. Zhang, P., Zhu, Y., Guo, Q., Li, J., Zhan, X., Yu, H., Xie, N., Tan, H., Lundholm, N., Garcia-Cuetos, L., et al. (2023) On the origin and evolution of RNA editing in metazoans. Cell Rep., 42(2):112112.

72. Denver, D.R., Morris, K., Kewalramani, A., Harris, K.E., Chow, A., Estes, S., Lynch, M. and Thomas, W.K. (2004) Abundance, distribution, and mutation rates of homopolymeric nucleotide runs in the genome of Caenorhabditis elegans. J. Mol. Evol., 58, 584–595.

73. Chen, N., Harris, T.W., Antoshechkin, I., Bastiani, C., Bieri, T., Blasiar, D., Bradnam, K., Canaran, P., Chan, J., Chen, C.-K., et al. (2005) WormBase: a comprehensive data resource for Caenorhabditis biology and genomics. Nucleic Acids Res., 33, D383–389.

74. Rosell, M., Rodríguez-Lumbreras, L.A. and Fernández-Recio, J. (2020) Modeling of Protein Complexes and Molecular Assemblies with pyDock. Methods Mol. Biol., 2165, 175–198.

75. Raitskin, O., Cho, D.S., Sperling, J., Nishikura, K. and Sperling, R. (2001) RNA editing activity is associated with splicing factors in lnRNP particles: The nuclear pre-mRNA processing machinery. Proc. Natl. Acad. Sci. U. S. A., 98, 6571–6576.

76. Hsiao, Y.-H.E., Bahn, J.H., Yang, Y., Lin, X., Tran, S., Yang, E.-W., Quinones-Valdez, G. and Xiao, X. (2018) RNA editing in nascent RNA affects pre-mRNA splicing. Genome Res., 28, 812–823.

77. Rodriguez, J., Menet, J.S. and Rosbash, M. (2012) Nascent-seq indicates widespread cotranscriptional RNA editing in Drosophila. Mol Cell, 47, 27–37.

78. Marcucci, R., Brindle, J., Paro, S., Casadio, A., Hempel, S., Morrice, N., Bisso, A., Keegan, L.P., Del Sal, G. and O’Connell, M.A. (2011) Pin1 and WWP2 regulate GluR2 Q/R site RNA editing by ADAR2 with opposing effects. EMBO J., 30, 4211–4222.

79. Nguyen Ba, A.N., Pogoutse, A., Provart, N. and Moses, A.M. (2009) NLStradamus: A simple Hidden Markov Model for nuclear localization signal prediction. BMC Bioinformatics, 10, 1–11.

80. Lin, J. rong and Hu, J. (2013) SeqNLS: nuclear localization signal prediction based on frequent pattern mining and linear motif scoring. PLoS One, 8(10):e76864.

81. Bernhofer, M., Goldberg, T., Wolf, S., Ahmed, M., Zaugg, J., Boden, M. and Rost, B. (2018) NLSdb-major update for database of nuclear localization signals and nuclear export signals. Nucleic Acids Res., 46, D503–D508.

82. Kosugi, S., Hasebe, M., Tomita, M. and Yanagawa, H. (2009) Systematic identification of cell cycle-dependent yeast nucleocytoplasmic shuttling proteins by prediction of composite motifs. Proc. Natl. Acad. Sci. U. S. A., 106, 10171–10176.

83. Shimozono, S., Tsutsui, H. and Miyawaki, A. (2009) Diffusion of large molecules into assembling nuclei revealed using an optical highlighting technique. Biophys. J., 97, 1288–1294.

