## Supplemental Data for "ADBP-1 regulates ADR-2 nuclear localization to control editing substrate selection"

**SUPPLEMENTARY DATA**

**Supplemental Figures**

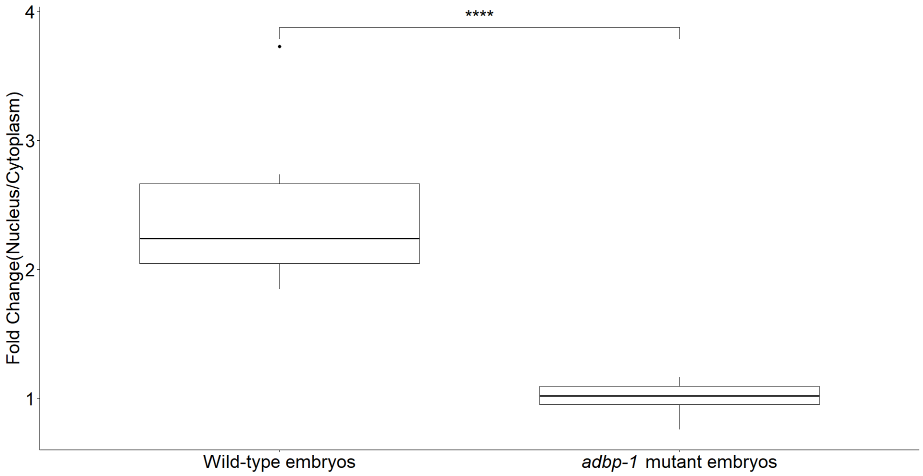


Supplemental Figure 1. Signal intensity quantification of ADR-2 subcellular localization from embryo immunostaining experiments. The Y-axis is the fold change expression of the mean gray value between the nucleus/cytoplasm after subtracting each fraction by the mean gray value of the background, which is calculated from micrographs from the immunostaining of ADR-2 in wild-type or *adbp-1* -/- embryos (Figure 1A). The same calculation was done for control on the *adr-2* mutant strain, and the signal was similar to the background. Background, cytoplasm fraction, and nucleoplasm fraction were taken for each cell quantified. At least ten cells were quantified for each strain from at least five different embryos. *** P-value < 0.001.

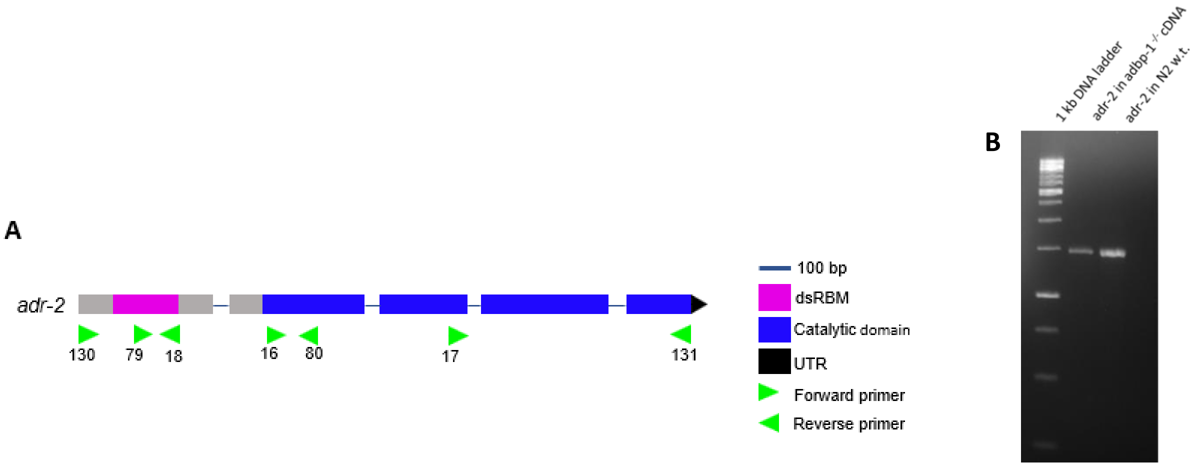


Supplemental Figure 2. Validation of *adr-2* gene in *adbp-1* mutant worms (A) Schematic view of the *adr-2* gene and the primers used to validate its sequence in *adbp-1* mutant worms. The gene is represented in its relative length. The double-strand RNA binding motif (dsRBM) region is shown in magenta, the deamination catalytic domain is shown in blue, and the UTR is shown in black. The primers that were used are represented as arrowheads in the relative portion of the gene. Arrowheads pointing to the right represent forward primers, while those pointing to the left represent reverse primers. The numbers indicated below belong to the primers' names as noted in Supp Table 1. (B) Electrophoresis gel of ADR-2 cDNA in *adbp-1* mutant worms. ADR-2 was amplified using primers AL_OBN_130 and AL_OBN_131. The expected fragment size is 1488 nt.

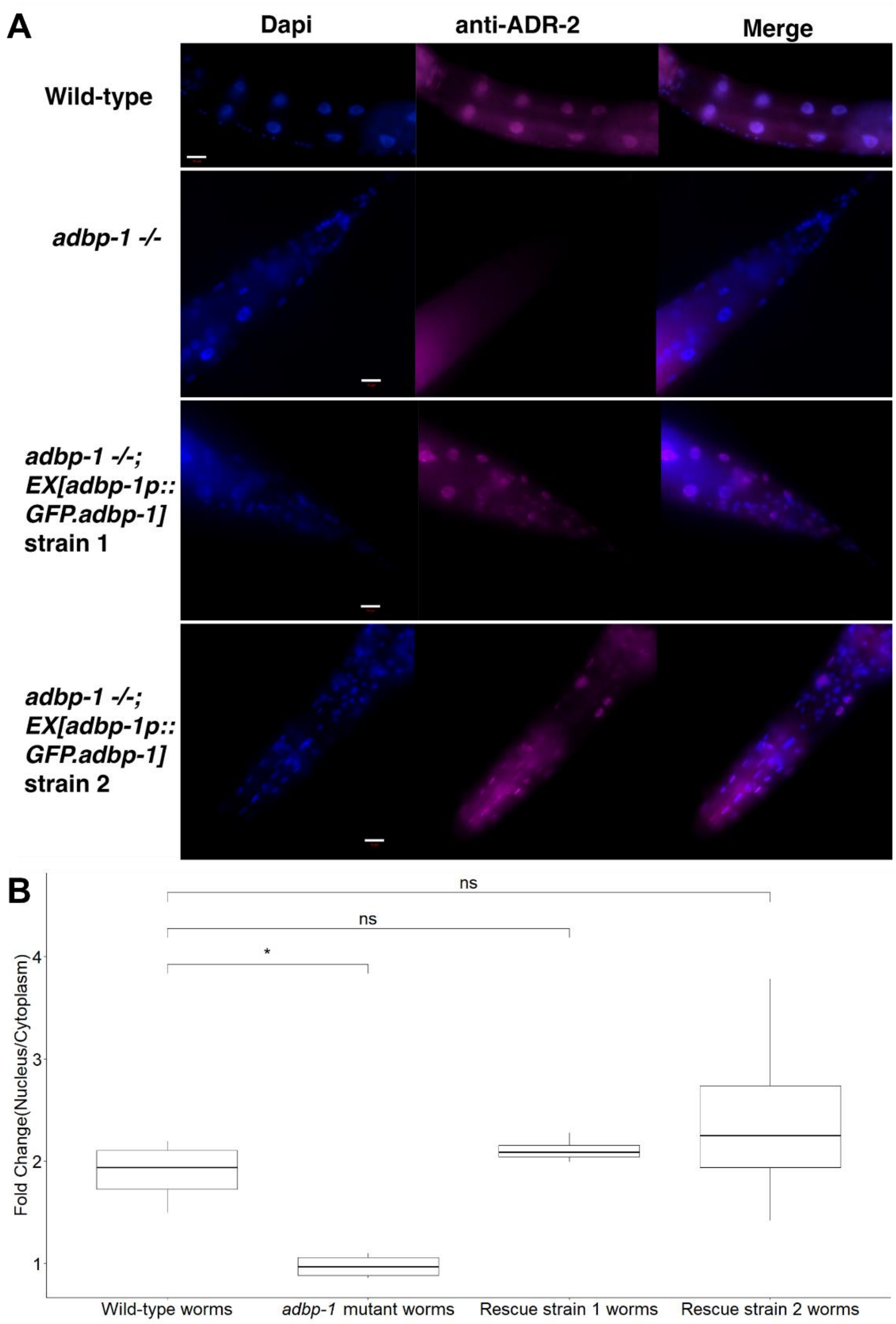


Supplemental Figure 3. Transgenic expression of ADBP-1 in the *adbp-1* mutant causes ADR-2 to be mainly nuclear as in wild-type. A. Representative immunofluorescence images of DNA (blue) and ADR-2 (magenta) of wild-type, *adbp-1(-/-),* and two strains expressing transgenic *adbp-1* under *adbp-1* promoter in *adbp-1 -/-* background: *adbp-1 (-/-)*; EX[*adbp-1*p::GFP.*adbp-1*] strains 1 and 2. Colocalization is shown as the overlap of the two images (merge). The scale bar in white is 10 µm. B. Quantification of ADR-2 signal in each subcellular localization from the immunostaining experiments in A in each strain. The Y-axis is the fold change expression of the mean gray value between the nucleus/cytoplasm after subtracting each fraction from the mean gray value of the background. Ns – non-significant, * P-value < 0.05.

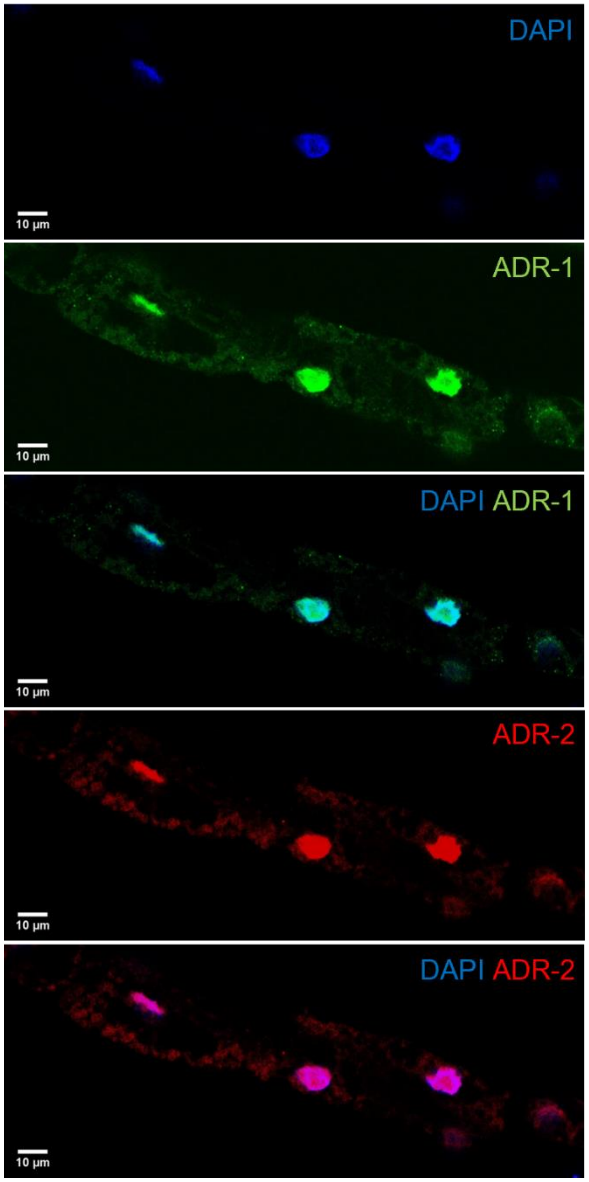


Supplemental Figure 4. ADR-1 and ADR-2 are localized in the nucleus. Representative confocal micrograph showing intestinal expression of ADR-1 and ADR-2 in adult *C. elegans*. Scale bar= 10µm. Both ADARs are concentrated in the intestinal nuclei, with dispersed signals in the cytoplasm. Intestine were isolated for the immunostaining.

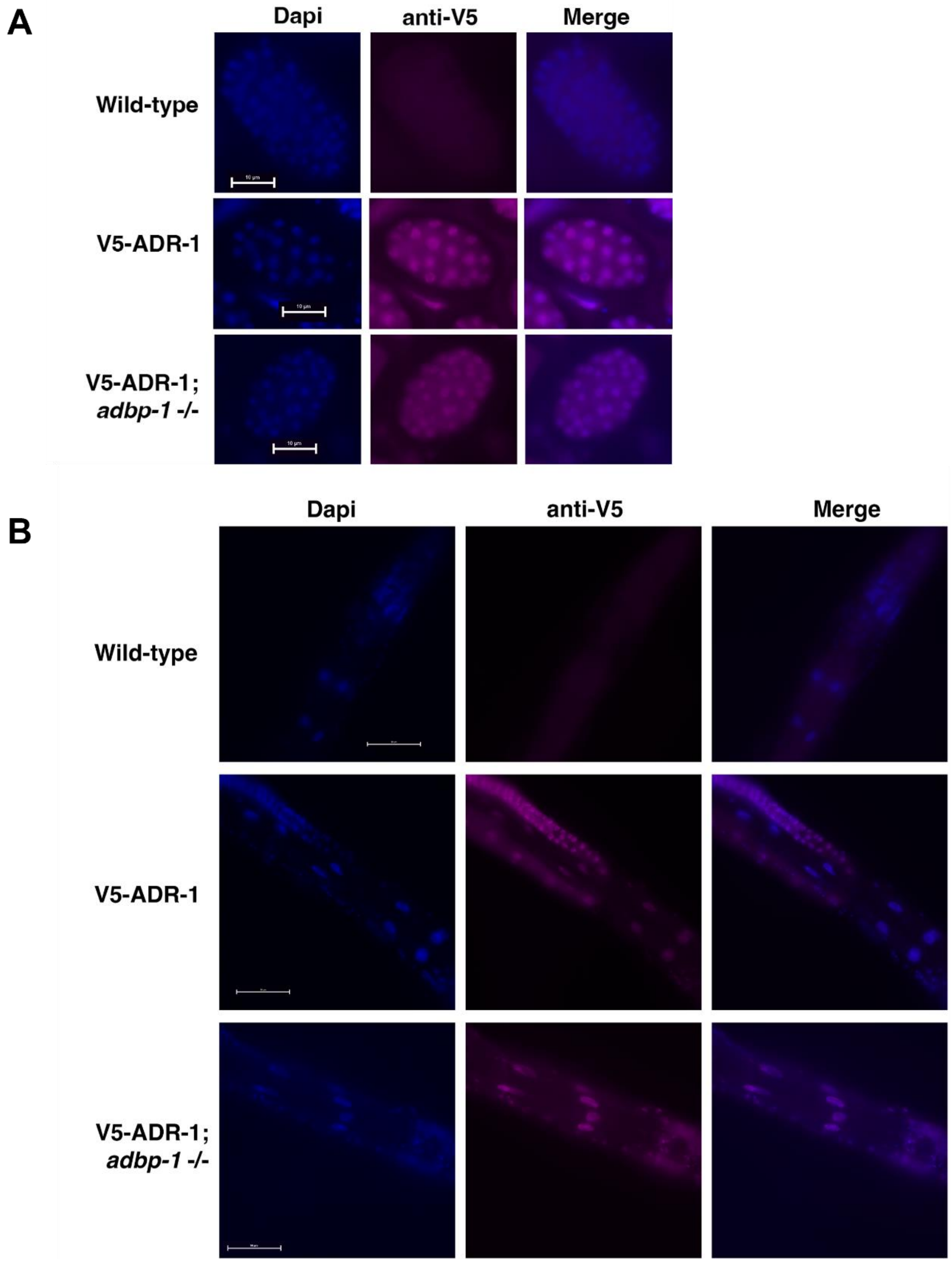


Supplemental Figure 5. ADR-1 subcellular expression does not change in *adbp-1 -/-*. Representative immunofluorescence images of DNA (blue) and V5-ADR-1 (magenta) in A. embryos and B. worm head of wild-type, worms expressing V5-ADR-1 and worms expressing V5-ADR-1 in *adbp-1 -/-* background. Antibody against the V5 epitope was used, which does not exist in wild-type worms. Colocalization is shown as the overlap of the two images (merge). The scale bar in A. is 10 µm, and in B. is 50 µm.

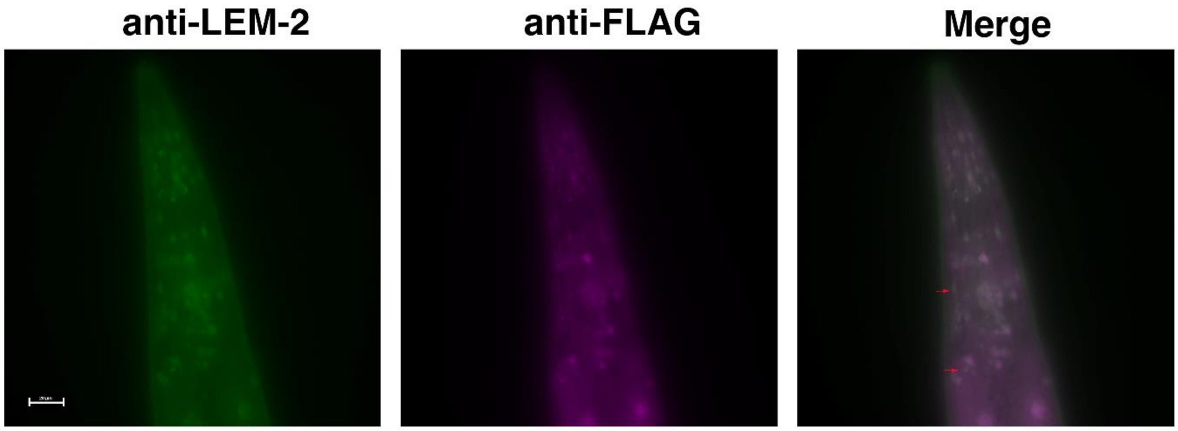


Supplemental Figure 6. LEM-2 is ubiquitously expressed, while ADR-2 is not. Representative immunofluorescence images of LEM-2 (green) and FLAG-ADR-2 (magenta) of FLAG-ADR-2 worms (HAH36 strain). Colocalization is shown as the overlap of the two images (merge). LEM-2 is expressed in the nuclear envelope. Red arrows point to cells that express LEM-2 but not ADR-2. Scale bar, 20 µm.

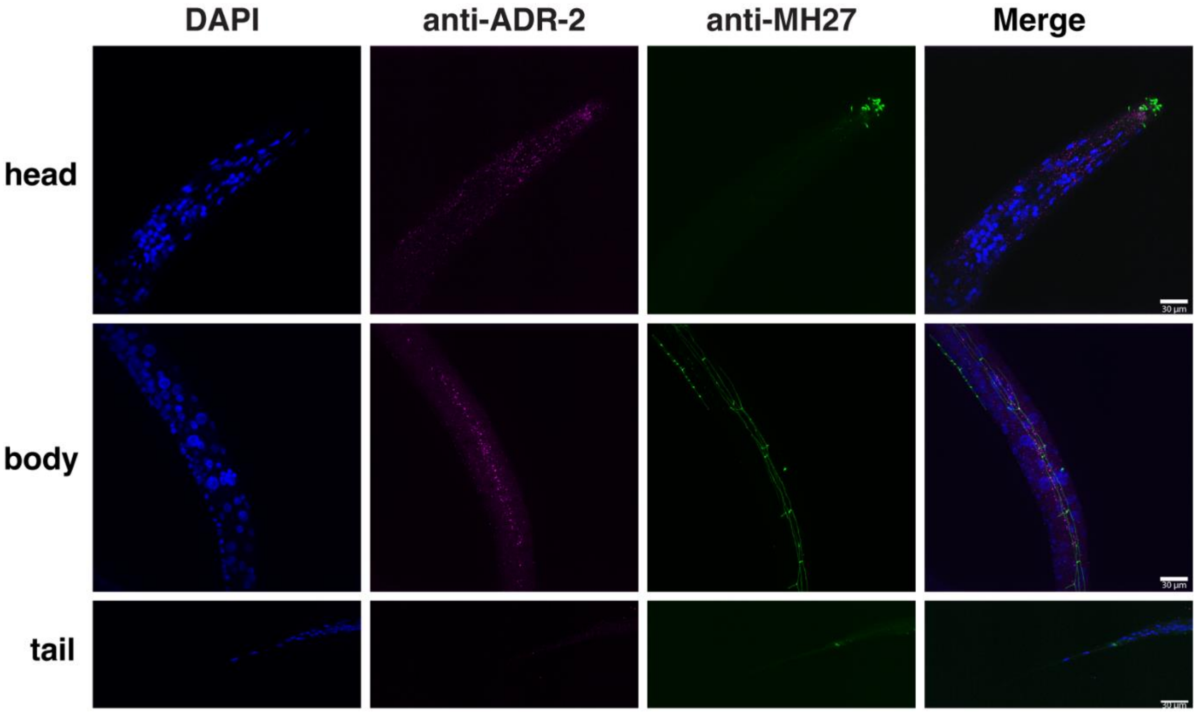


Supplemental Figure 7. Localization of ADR-2 in the *adbp-1(-/-)* hermaphrodite head, body and tail. Representative immunofluorescence images of DNA (blue), ADR-2 (magenta) and MH27 (green) from head (top), body (middle) and tail (bottom) of *adbp-1(-/-)*. Colocalization is shown as the overlap of the three images (merge). Scale bar, 30 µm

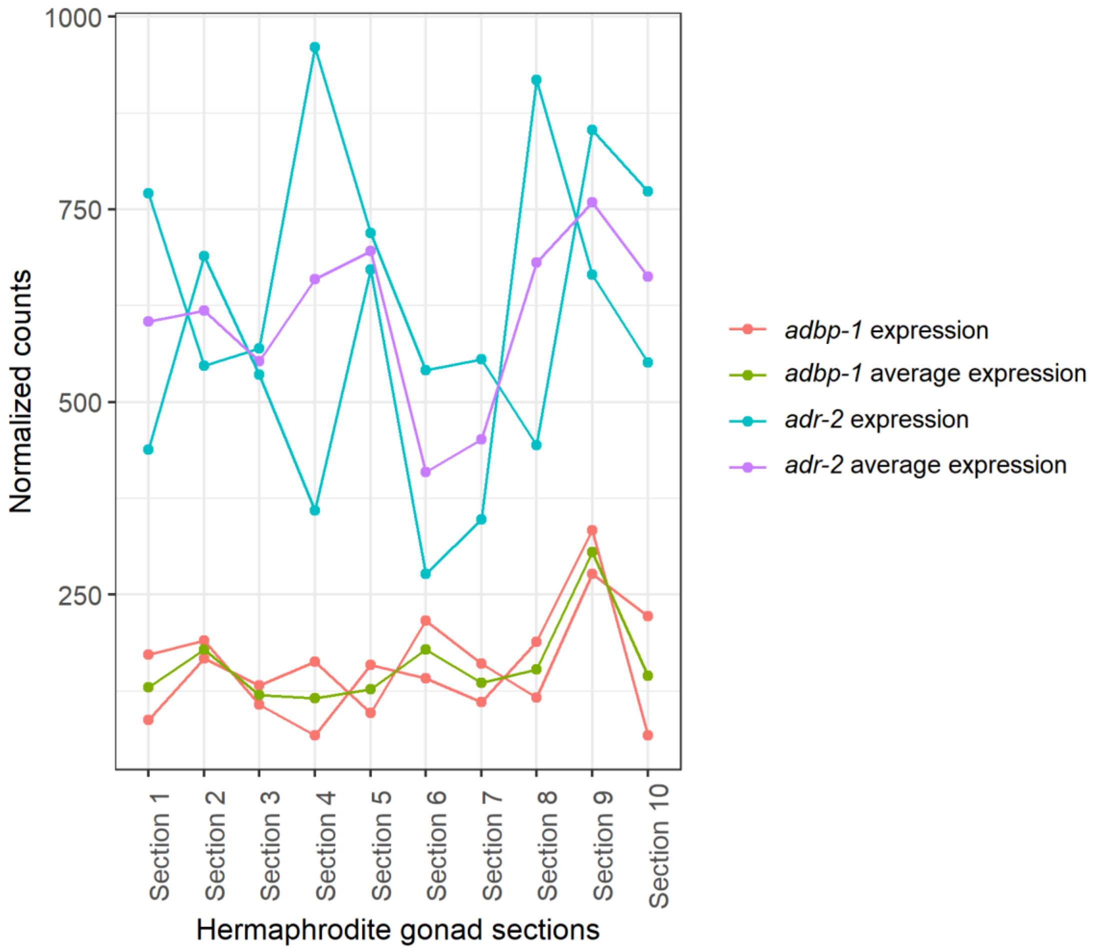


Supplemental Figure 8. *adr-2* and *adbp-1* mRNA levels in hermaphrodite's gonads. The plot shows *adr-2* and *adbp-1* gene expression analysis from Tzur et al., 2018 (1), in which single-cell RNA-seq was performed on dissected gonad section.The blue lines represent *adr-2* expression in two different biological replicas, while the purple line is the average *adr-2* expression. The orange lines represent *adbp-1* expression in two different biological replicas, while the purple line is the average *adbp-1* expression. Each gonads section represents a different development stage during oogenesis.

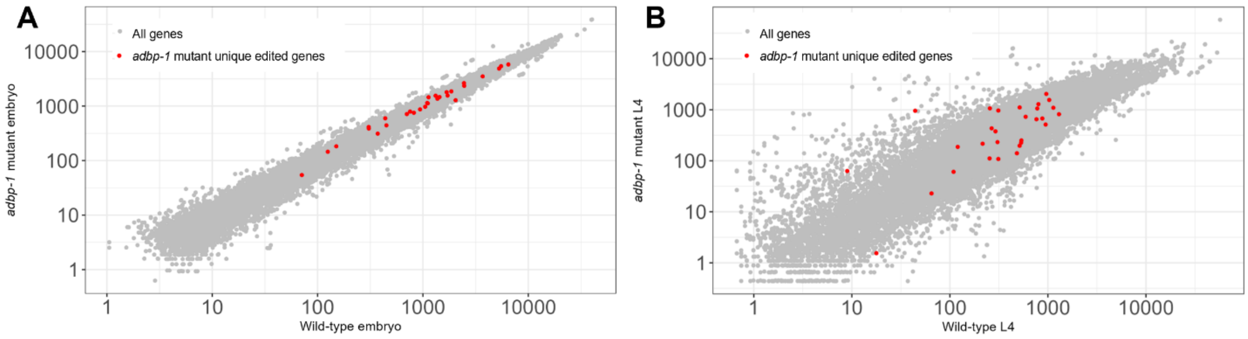


Supplemental Figure 9. Genes that are edited only in *adbp-1* mutant worms are highly expressed. Log scale plots represent the genes expressed in wild-type worms versus *adbp-1* mutant worms in gray in the embryo (A) and the L4 stage (B). Grey dots represent the average counts of the genes across biological replicas produced by DESeq2 (2). Unique edited genes to *adbp-1* mutant worms found in the computational pipeline searching for new editing sites are highlighted in red.

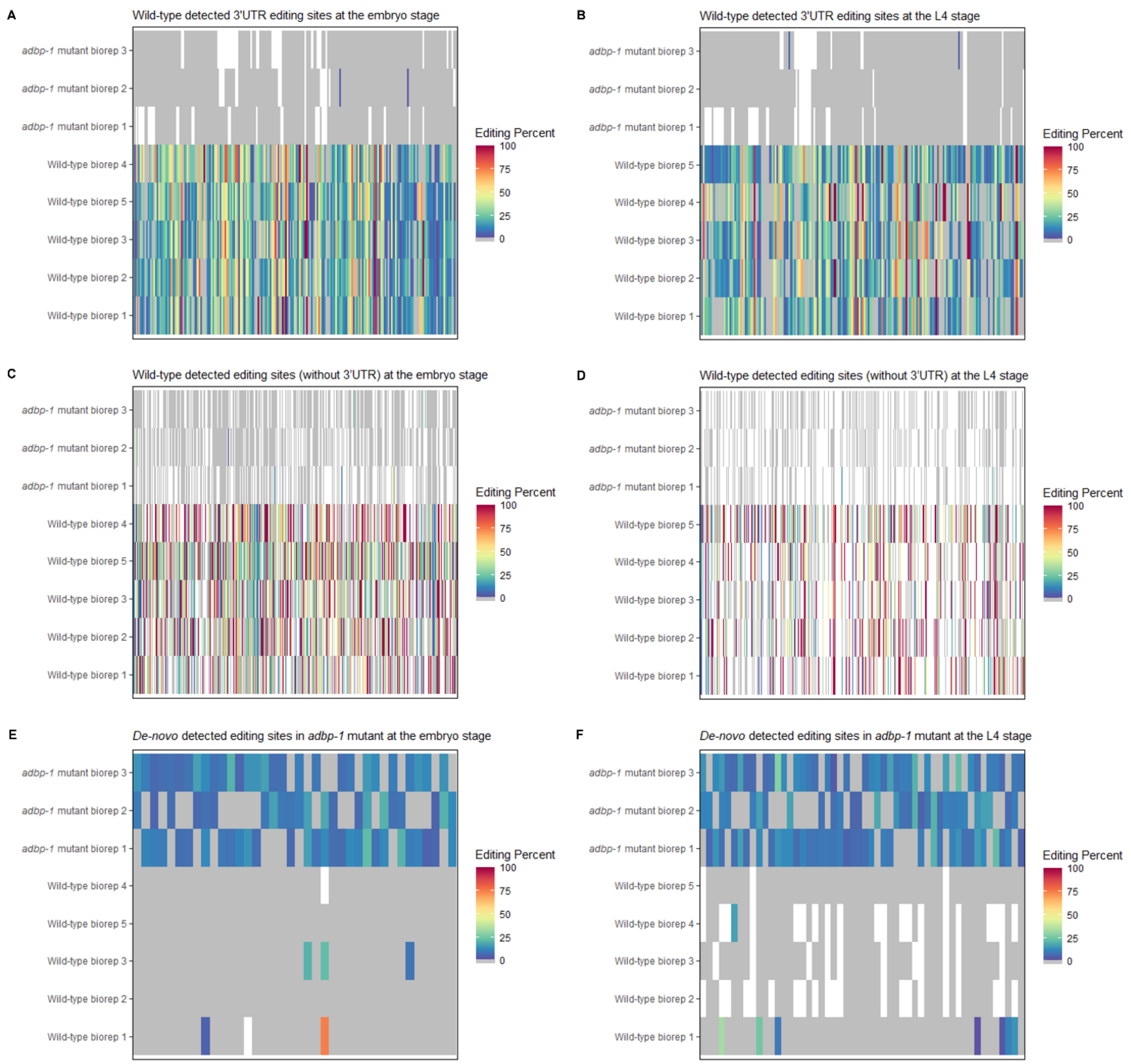


Supplemental Figure 10. Illustration of editing percentage in detected editing sites. The heatmaps illustrate the editing percentage values for various sites identified as edited in prior transcriptome-wide investigations (3, 4) and our novel *de-novo* editing site search. The heatmaps depict these editing percentage values across all samples utilized in our research. The analysis covers wild-type (A-D) and *adbp-1* mutant worms (E-F). In the heatmaps, areas without any expression are depicted in white. In contrast, sites that display expression but no editing (with an editing percentage of zero) are shown in gray. The scale ranges from blue to red, representing editing percentages from above zero to 100.

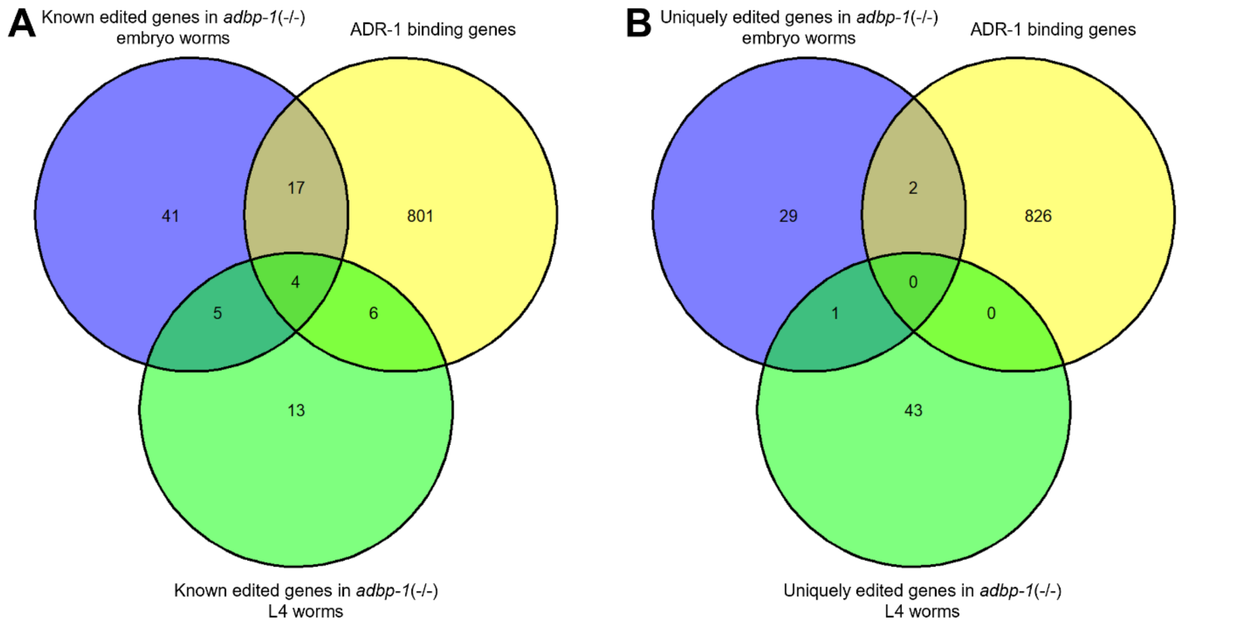


Supplemental Figure 11. Edited genes in *adbp-1* mutant worms significantly overlap with ADR-1 binding genes. The Venn diagrams represent (A) The intersection between editing sites found in transcriptome-wide studies (3, 4) (known edited sites) in *adbp-1* mutant worms at the embryo and L4 stages and ADR-1 binding genes. 21 known edited genes in *adbp-1* mutant worms at the embryo were found to be ADR-1 binding genes (P-value = 5.52e-15). 10 known edited genes in *adbp-1* mutant worms at the L4 stage were found to be ADR-1 binding genes (P-value = 2.61e-09). (B) The intersection between uniquely edited genes in *adbp-1* mutant worms at the embryo and the L4 stage and ADR-1 binding genes. Two uniquely edited genes in the *adbp-1* mutant worms at the embryo stage were found to be ADR-1 binding genes; however, the P-value is not significant. None of the uniquely edited genes in the *adbp-1* mutant worms at the L4 stage were found to be an ADR-1 binding gene. P-values were determined by a hypergeometric test.

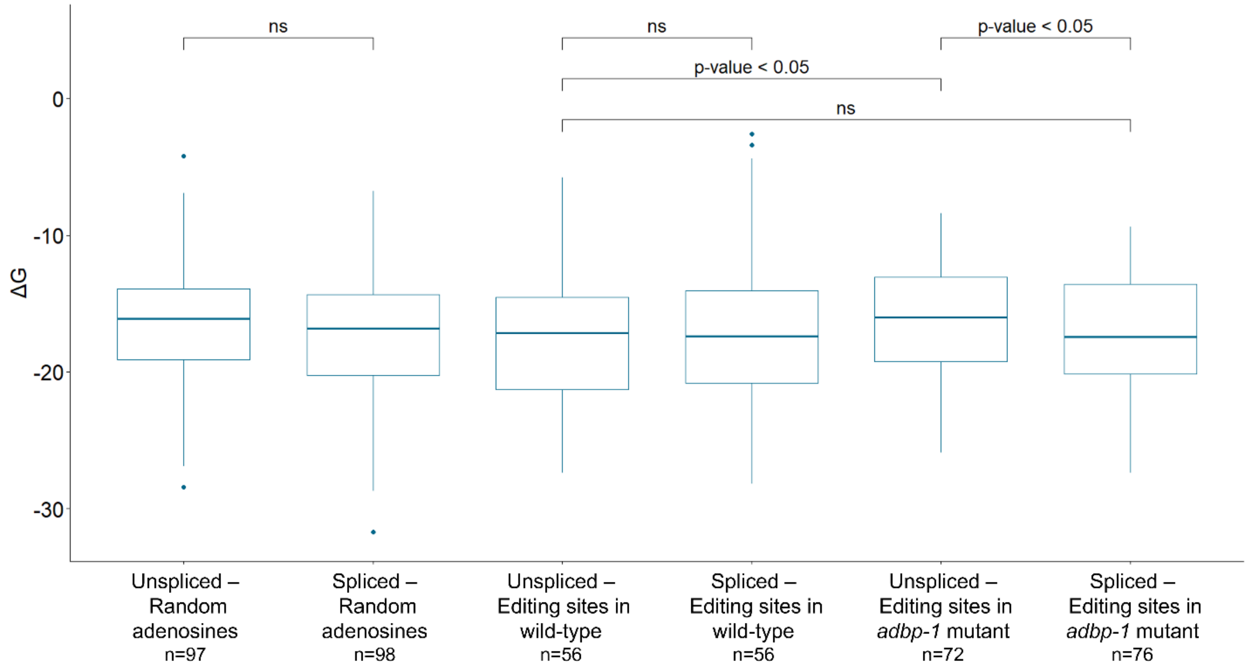


Supplemental Figure 12. Free energy of edited sequences secondary structure. The box plots illustrate the average minimum free energy of 101-nucleotide-long secondary structures surrounding randomly selected adenosine positions and those identified as edited. These secondary structures are examined in their unspliced and spliced forms within wild-type and *adbp-1* mutant worms. "ns" stands for "non-significant" values. P-values were calculated using the Welch two-sample T-test.

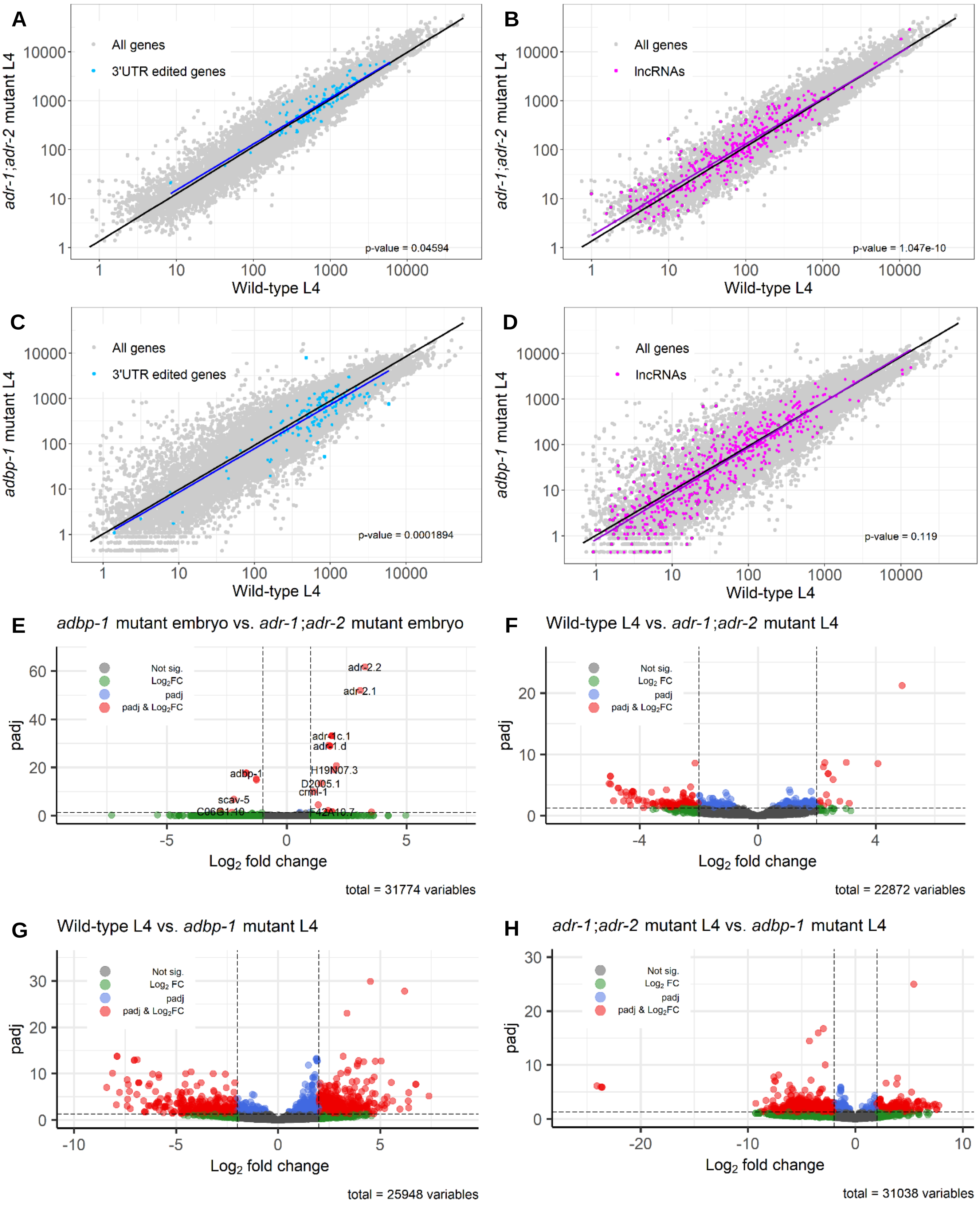
A
All genes
3'UTR edited genes
adr-1;adr-2 mutant L4
Wild-type L4
p-value = 0.04594
B
All genes
lncRNAs
adr-1;adr-2 mutant L4
Wild-type L4
p-value = 1.047e-10
C
All genes
3'UTR edited genes
adbp-1 mutant L4
Wild-type L4
p-value = 0.0001894
D
All genes
lncRNAs
adbp-1 mutant L4
Wild-type L4
p-value = 0.119
E
adbp-1 mutant embryo vs. adr-1;adr-2 mutant embryo
Not sig.
Log2 FC
padj
padj & Log2FC
adr-2.2
adr-2.1
adr-1c.1
adr-1.d
H19N07.3
adbp-1
D2005.1
cml-1
scav-5
C06G1.10
F42A10.7
padj
Log2 fold change
total = 31774 variables
F
Wild-type L4 vs. adr-1;adr-2 mutant L4
Not sig.
Log2 FC
padj
padj & Log2FC
padj
Log2 fold change
total = 22872 variables
G
Wild-type L4 vs. adbp-1 mutant L4
Not sig.
Log2 FC
padj
padj & Log2FC
padj
Log2 fold change
total = 25948 variables
H
adr-1;adr-2 mutant L4 vs. adbp-1 mutant L4
Not sig.
Log2 FC
padj
padj & Log2FC
padj
Log2 fold change
total = 31038 variables

Supplemental Figure 13. Low editing levels affect genes expression at the L4 stage. (A-D) The genes expressed in wild-type worms versus *adbp-1* mutant worms and the genes expressed in wild-type worms versus ADAR mutant worms at the L4 stage are represented in log scale plots. Each dot represents an expressed gene. Grey dots represent all the genes, blue dots represent edited genes at their 3'UTR, and purple dots represent lncRNAs. The black line is a regression line for all genes, the blue line is the regression line for genes that are edited at their 3'UTR, and the purple line is the regression line for the lncRNAs. (E-H) The volcano plots depict the log2 fold change versus -log10(P-adjusted) between the genes expressed in wild-type worms to *adr-1;adr-2* mutant worms, and wild-type worms to *adbp-1* mutant worms at the embryo. Non-significant genes are colored in grey. (E) Differentially expressed genes, which adhere to the following criteria: |log2FoldChange| > 2 and P-adjusted < 0.05, are highlighted in red. Genes with only |log2FoldChange| > 2 are colored green, and genes with only P-adjusted < 0.05 are colored blue (F-H).

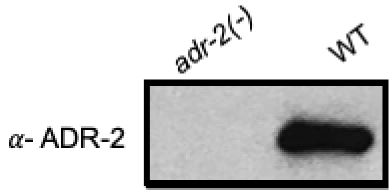


Supplemental Figure 14. A western blot of ADR-2 IPs was submitted to mass spectrometry.

**Supplemental Tables**

| Primer name | Sequence | Description |
|---|---|---|
| AL_OBN_130 | ATGTCCGTCGAAGAAGGTATG | Forward primer for 5' of *adr-2* |
| AL_OBN_131 | TTAATTTATAGTAAACATTTGAAATTCTCGTG | Reverse primer for 3' of *adr-2* |
| AL_NS_16 | CAAGGGATTACGCGGAGAC | Forward primer for *adr-2* Sanger sequencing validation |
| AL_NS_17 | TACTGGGTGTCCAAGGAG | Forward primer for *adr-2* Sanger sequencing validation |
| AL_OBN_79 | TCTATAACAGAAACCAACCG | Forward primer for *adr-2* Sanger sequencing validation |
| AL_OBN_80 | CTCTTCTTGCGAGAATTTC | Reverse primer for *adr-2* Sanger sequencing validation |
| AL_NS_18 | CACGACACCTTTGCCTCTTTC | Reverse primer for *adr-2* Sanger sequencing validation |
| AL_NS_19 | GATGGAAATACAGAAGTTG | Forward primer for DNA of *adr-2* before the translation start site |
| AL_NS_20 | CGTTCACTAGTCGATGTTGCTCTATTG | Forward primer for DNA *adr-2* Sanger sequencing validation |
| AL_OBN_184 | AAAAGCTGAAGAAACAGGAC | Forward primer for 5' of *adbp-1:gfp* "L" plasmid |
| AL_OBN_185 | TTAACATCACCATCTAATTCAAC | Reverse primer for 3' of *adbp-1:gfp* "L" plasmid |

Supplemental Table 1. Primers were used in this study.

| | Wild-type Embryo biorep 1 | Wild-type Embryo biorep 2 | Wild-type Embryo biorep 3 | Wild-type Embryo biorep 4 | Wild-type Embryo biorep 5 |
|---|---|---|---|---|---|
| 2 samples | 8.4% | 7.8% | 10.2% | 10.9% | 12.7% |
| 3 samples | 5.8% | 6.1% | 7.2% | 7.1% | 7.2% |
| 4 samples | 6.3% | 6.9% | 6.8% | 6.8% | 6.3% |
| 5 samples | 8.5% | 8.3% | 7.4% | 9.3% | 6.6% |
| | *adbp-1* mutant Embryo biorep 1 | *adbp-1* mutant Embryo biorep 2 | *adbp-1* mutant Embryo biorep 3 | | |
| 2 samples | 2.1% | 2.5% | 2.6% | | |
| 3 samples | 0.2% | 0.2% | 0.2% | | |
| | Wild-type L4 biorep 1 | Wild-type L4 biorep 2 | Wild-type L4 biorep 3 | Wild-type L4 biorep 4 | Wild-type L4 biorep 5 |
| 2 samples | 4.6% | 5.6% | 5.2% | 3.1% | 9.4% |
| 3 samples | 3.1% | 4.4% | 4.4% | 1.4% | 6.2% |
| 4 samples | 3.3% | 4.6% | 4.3% | 1.2% | 5% |
| 5 samples | 2.6% | 3.6% | 3.2% | 2.2% | 3.7% |
| | *adbp-1* mutant L4 biorep 1 | *adbp-1* mutant L4 biorep 2 | *adbp-1* mutant L4 biorep 3 | | |
| 2 samples | 2.8% | 2.6% | 2.2% | | |
| 3 samples | 0.2% | 0.2% | 0.2% | | |

Supplemental Table 2. Editing sites reproducibility across biological replicas of wild-type and *adbp-1* mutant worms. The table shows, for each biological replica, the percentage of editing sites that appear in two samples and more in wild-type and *adbp-1* mutant worms at the embryo and L4 developmental stages out of the overall editing site detected in the replica.

Supplemental Table 3. List of sites found in the *adbp-1* mutant strain that do not appear in wild-type worms. New_editing_sites_in_adbp-1_mutant_worms.xlsx

Supplemental Table 4. Differential expression of genes between wild-type, *adbp-1* mutant and ADR-2 mutant worms. Differential_expression_analysis_results.xlsx

| **Protein name** | **Gene locus** | **Size** | ***p*-value (WT vs *adr-2(-)*)** | **Unique peptides in WT** | **Unique peptides in *adr-2(-)*** |
|---|---|---|---|---|---|
| Double-stranded RNA-specific adenosine deaminase | *adr-2* | 55 kDa | <0.00010 | 32 | 0 |
| Cluster of Adenosine Deaminase acting on RNA | *adr-1* | 110 kDa | <0.00010 | 32 | 0 |
| ADR-2 binding protein | *adbp-1* | 24 kDa | <0.00010 | 21 | 0 |
| Importin subunit alpha-3 | *ima-3* | 56 kDa | 0.034 | 3 | 0 |
| Importin beta family | *Imb-3* | 122 kDa | 0.04 | 7 | 1 |

Supplemental Table 5. ADR-2 interacting proteins identified using immunoprecipitation and mass spectrometry. The identified ADR-2 interacting proteins are listed with protein name, gene locus, and size in the first three columns. Statistical significance for the enrichment of unique peptides in IPs from wild-type (WT) and *adr-2*(-) worms was calculated using Fisher's t-test in Scaffold and the calculated p values are listed in column 4. The number of unique peptides corresponding each listed protein in IPs are listed in the next two columns. Two biological replicates were performed for each strain.

| | Model 1 | Model 2 | Model 3 | Model 4 | Model 5 | **Average** |
|---|---|---|---|---|---|---|
| ADR2-ADBP1_full | -109.409 | -112.889 | -126.358 | -108.280 | -128.334 | **-116.6** |
| ADR2-ADBP1_stop | -67.403 | -93.138 | -75.452 | -108.056 | -104.019 | **-89.6** |

Supplemental Table 6. pyDock Interface total energy for the full and mutated ADBP1. Despite both interfaces are stable, the full complex has a significantly higher interface energy average, suggesting a more stable interface.

**REFERENCES**


1. Yonatan B. Tzur, Eitan Winter, Jinmin Gao, Tamar Hashimshony, Itai Yanai, and MPC (2018) Spatiotemporal Gene Expression Analysis of the Caenorhabditis elegans Germline Uncovers a Syncytial Expression Switch. *Genetics*, **210**, 587–605.

2. Anders,S. and Huber,W. (2010) Differential expression analysis for sequence count data. *Genome Biol.*, **11**, R106.

3. Goldstein,B., Agranat-Tamir,L., Light,D., Zgayer,O.B.N., Fishman,A. and Lamm,A.T. (2017) A-to-I RNA editing promotes developmental stage-specific gene and lncRNA expression. *Genome Res.*, **27**, 462–470.

4. Ganem,N.S., Ben-Asher,N., Manning,A.C., Deffit,S.N., Washburn,M.C., Wheeler,E.C., Yeo,G.W., Zgayer,O.B.N., Mantsur,E., Hundley,H.A., *et al.* (2019) Disruption in A-to-I Editing Levels Affects C. elegans Development More Than a Complete Lack of Editing. *Cell Rep.*, **27**, 1244-1253.e4.
